# Condensin I^DC^ has a Functional ATPase That is Required for X-Chromosome Dosage Compensation in *C. elegans*

**DOI:** 10.1101/2025.02.05.636654

**Authors:** Bahaar Chawla, Suchi Jatia, Dillon E. Sloan, Gyorgyi Csankovszki

**Affiliations:** Department of Molecular, Cellular, and Developmental Biology, University of Michigan, Ann Arbor, MI 48109

## Abstract

Dosage compensation (DC) in *C. elegans* utilizes a condensin complex that resembles mitotic condensins, but differs by one subunit, DPY-27. DPY-27 replaces SMC-4, one of the Structural Maintenance of Chromosome (SMC) proteins that is responsible for hydrolyzing ATP, required for condensation of DNA and other mitotic condensin functions. To understand if the ATPase function is required in DC, we first demonstrated that DPY-27 is capable of hydrolyzing ATP *in vitro*. Then, we used CRISPR/Cas9-mediated genome editing to generate an ATPase mutation in *dpy-27* and demonstrated that this mutation results in a loss of DC. Specifically, we found that without ATPase function, DPY-27 containing condensin I^DC^ has reduced capacity to bind DNA, condense the X chromosomes, and facilitate H4K20me1 enrichment on the X-chromosomes. Our results suggest that condensin I^DC^, like mitotic condensins, uses ATP hydrolysis to perform its functions, making *C. elegans* DC a model for how activities attributed to mitotic condensins can be used to regulate gene expression.

## Introduction

Our understanding of chromatin structure has evolved greatly through the history of genetics. Beginning with Flemming’s dark-stained bodies [1] to our modern understanding of the multiple levels of chromatin compaction [2–7], we recognize that chromatin is a highly dynamic and constantly changes through the cell cycle [8,9], with gene expression changes [10,11], or tissue specification [12].

One of the protein complexes capable of compacting chromatin is condensin [13,14]. Condensins are five-subunit protein complexes, found throughout bacteria and eukaryotes [13,15–17]. The core of condensin is made up of the two Structural Maintenance of Chromosome (SMC) proteins and a kleisin [18–22]. In bacterial condensins, there is a single SMC protein that functions as a homodimer [15,23]. However, in eukaryotes, there are multiple SMC proteins [13,18,24–26]. Condensin has the heterodimeric pair of SMC2 and SMC4 [13,24]. Another pair of SMC proteins are found in the very similar and related complex, cohesin that hold sister chromatids together [18,27]. Cohesin also contains a kleisin [17,20,28,29]. The final pair of eukaryotic SMC proteins are found in the SMC5/6 complex that is involved in DNA repair, transposon silencing, and other functions distinct from chromosome compaction [25,30–32]. The final two subunits of a condensin are the HEAT-proteins Associated With Kleisins (HAWKs) which use their HEAT domains to interact with the kleisin and SMC proteins [33–37].

In *Caenorhabditis elegans*, the microscopic nematode, like many other eukaryotes, there are two distinct condensin complexes active during mitosis, condensin I and II [16,38–44]. Condensin I and II share the same SMC proteins MIX-1 (the common name for SMC2 in *C. elegans*) and SMC-4, but different kleisins and HAWKs [16,41,44]. Condensin I does not localize to DNA until after nuclear envelope breakdown [42,45], and then coats the chromosomes in a pattern distinct from condensin II [16,43]. Meanwhile, condensin II is present in the nucleus during prophase [45], just like in other eukaryotes [42]. During anaphase, *C. elegans* condensin I localizes to the spindle midzone where it helps resolve chromatin bridges [46]. Studies in other organisms revealed that many of the mitotic processes involving condensin require ATP, and the two SMC proteins in condensin work together as an ATPase [26,39,47,48]. When this ATPase function is compromised, condensins are unable to compact DNA [48]. Condensin’s ability to hydrolyze ATP was originally demonstrated in *P. furiosus*. Comparisons between the ABC (ATPase Binding Cassettes) domains in Rad50 [49,50] and the head domains of condensin showed a high degree of similarity and were used as the basis to test if SMC proteins were able to bind and hydrolyze ATP [15]. Moreover, it was found that mutating the essential glutamate to a glutamine (EQ) in the Walker B domain abolished ATPase capability [15]. This residue has been shown to be highly conserved, from bacteria to yeast to humans, and generation of the EQ mutation is sufficient to abolish ATPase capability in all species tested [26,48,51]. Moreover, loss of ATPase capability reduced DNA binding *in vitro* [51], prevented DNA compaction, and resulted in a loss of cell viability in yeast [47,48]. Interestingly, research now shows that condensins are also found in interphase, where they may play a gene regulatory role [52–54]. However, it is not known whether their ATPase activity is required for these interphase functions.

Unlike other eukaryotes, *C. elegans* contain an extra condensin [16,44] (Figure 1A). This extra condensin is known as condensin I^DC^ because it functions specifically in dosage compensation (DC). *C. elegans* has two sexes, a male and a hermaphrodite, which differ by their number of X chromosomes, X and XX respectively [55]. This dosage imbalance between the sexes is ameliorated by the process of dosage compensation [56,57]. The Dosage Compensation Complex (DCC) is a ten-subunit protein complex. The first five are the proteins making up condensin I^DC^; the two SMC proteins MIX-1 and DPY-27 (a paralogue of SMC-4), and the same kleisin and HAWKS found in condensin I: DPY-26, DPY-28, and CAPG-1 [16,44,58–62]. The other members of the complex include SDC-1, SDC-2, and SDC-3, named for their roles in both Sex Determination and Dosage Compensation [63–65]. The final two members are DPY-21, a H4K20me2 demethylase [66,67], and DPY-30, which is also a member of the MLL/COMPASS complex but its exact role in DC is still under investigation [68,69]. Once localized to X chromosomes [70,71], the DCC reduces gene expression of each hermaphrodite X by half so that the total gene output equals the single male X [57,72].

**Figure 1.**
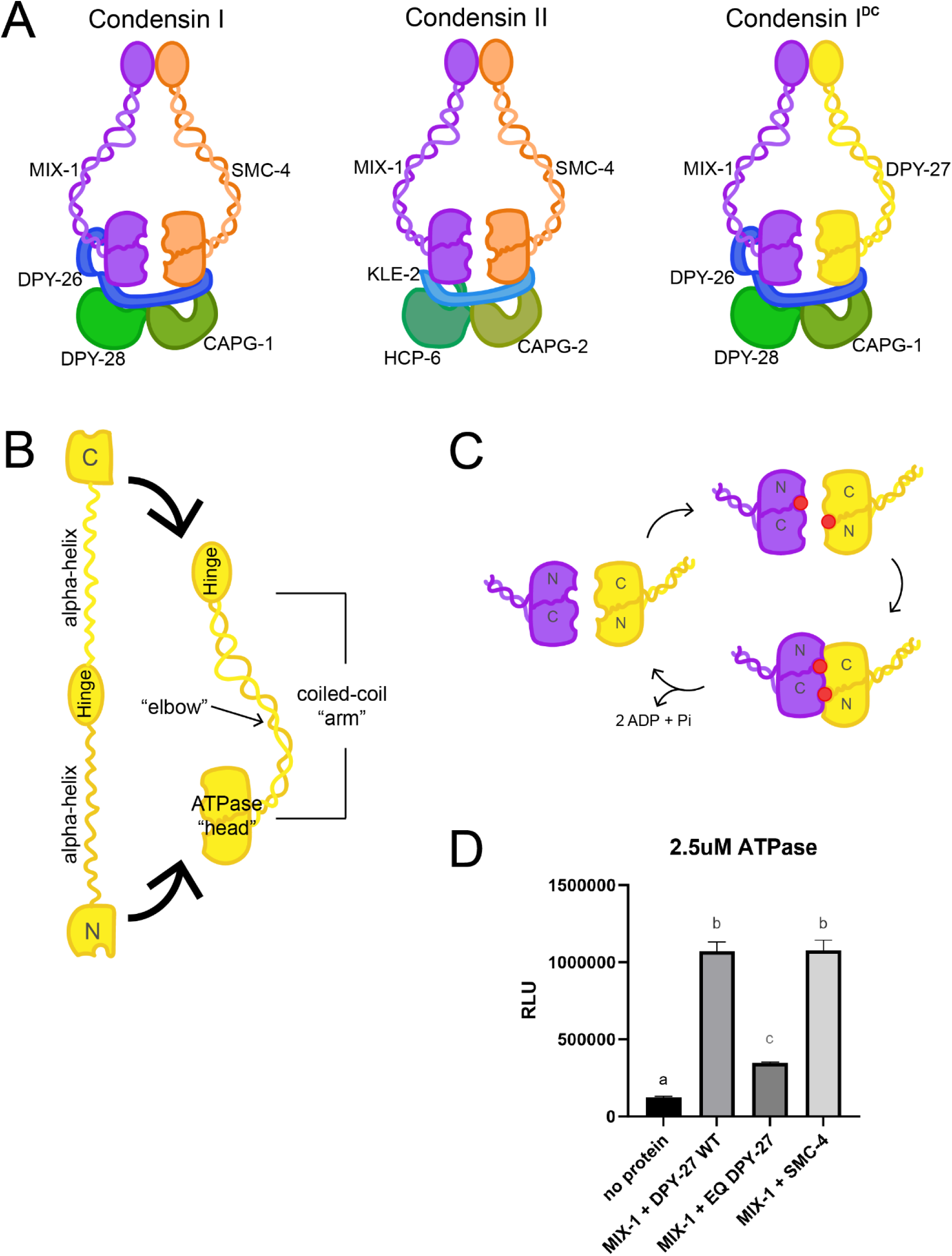
Condensins in *C. elegans* and ATPase function of Condensin I^DC^. **(A)** *C. elegans* has three condensin complexes. Condensin I and II function in mitosis and meiosis. Condensin I^DC^ functions uniquely in DC, and is identical to condensin I, except that the DC specific SMC protein DPY-27 replaces SMC-4. The kleisins are in shades of blue, and the HAWKs are in shades of green. **(B)** SMC proteins are made up of two globular domains separated by a pair of alpha-helices and a “hinge” domain. Once the hinge folds, the alpha-helices become a coiled-coil that brings together the N and C terminal domains into the “ATPase head”. The coiled-coil is sometimes referred to as the “arm” with an “elbow” where bending can occur. **(C)** The current proposed mechanism of how SMC proteins hydrolyze ATP is shown. Here, the walker A in the N-terminal portion of the head binds an ATP and once the two heads come together in space, the opposing head’s walker B (located in the C-terminal portion) removes the phosphate group. The two heads then return to their open confirmation. **(D)** The rate of ATP hydrolysis was assayed for the different SMC “ATPase head” protein heterodimers. Relative luminescence units, RLU, is directly proportional to the amount of ADP produced. MIX-1 and DPY-27 WT together were capable of hydrolyzing ATP at a rate similar to the mitotic SMC head pair, while the EQ DPY-27 mutant head with MIX-1 severely reduced ATP hydrolysis. 3 replicates of each reaction were run and a one-way ANOVA with multiple comparisons was used to determine statistical significance. Letters represent multiple comparison p values, with different letters indicating statistically significant differences, and any repeated letter demonstrating no significance. See Supplemental Table 4 for all p-values.

The existence of DPY-27, a subunit unique to condensin I^DC^, allows for the dissection of condensin-mediated gene regulation on a wide scale. While condensins have been shown to impact gene expression during interphase [52–54], such studies are limited due to the challenge of decoupling condensin’s mitotic role from their gene regulatory role. DPY-27 in *C. elegans* allows studies on how condensin I^DC^ directly impacts gene regulation on a chromosome-wide scale, without interrupting cell division and growth. One key question is whether condensin I^DC^ can hydrolyze ATP like other SMC protein complexes [15,26,47,50]. Previous research using *C. elegans* has demonstrated the existence of Topologically-Associated Domains (TADs) specifically in hermaphrodite X chromosomes, which are dependent on the DCC [73,74]. TADs in other organisms are generated by cohesin [75]. Cohesin resembles condensin, containing two SMC proteins and a kleisin, but differs from condensin as cohesin contains only one HAWK [18]. TADs formed by cohesin are dependent on ATP hydrolysis by the SMC proteins in the complex [76].

When comparing homology between SMC proteins, DPY-27 clearly resembles other SMC proteins (Supplementary Figure 1). SMC proteins have a stereotypical tertiary structure where the N and C-termini are large globular domains separated by two α-helices and a hinge [15,18,21,51]. The hinge folds and the helices become a coiled-coil, bringing the N and C-termini together (Figure 1B). The resulting “ATPase head” has been shown to be capable of hydrolyzing ATP in many different organisms, including bacteria, humans, and yeast [15,39,48]. The proposed mechanism involves two SMC protein ATPase heads which work together to hydrolyze two ATP molecules simultaneously [15,26]. Each head binds an ATP molecule, using the Walker A domain and the Walker B of other head performs the hydrolysis (Figure 1C). However, it is not known if DPY-27 is capable of hydrolyzing ATP. One hypothesis that would explain the difference between SMC-4 and DPY-27 is that SMC-4 has ATPase function as a mitotic condensin subunit, but DPY-27 does not and is only a scaffold protein. Previous research with an “ATPase-dead” mutant protein *in vivo* showed no incorporation of the mutant DPY-27 into condensin I^DC^ and instead degradation of the mutant DPY-27 species over time [77]. However, this “ATPase-dead” mutant was a transgenic allele under the control of a heat-shock promoter in a genetic background that also contained a wild type *dpy-27* allele.

Here we demonstrate *in vitro* that DPY-27 is capable of hydrolyzing ATP and the EQ mutation abolishes this function. We also generated a worm with the “ATPase-dead” EQ mutation, analogous to mutations generated in many other species [15,26,47], in the endogenous *dpy-27* locus. Surprisingly, this mutant did not phenocopy the transgenic EQ mutant in [77], leading us to carefully analyze mutant phenotypes. We found that the EQ DPY-27 mutant protein colocalizes with other subunits of condensin I^DC^ but mutant worms do not have proper dosage compensation. We also demonstrated that DPY-27 EQ mutant protein has varied DNA binding ability. Finally, we observed that loss of the ATPase function affects other DC processes. Taken together, our results suggest that condensin I^DC^ is a loop-extruding SMC complex like other condensins, and how it regulates the X chromosomes in *C. elegans* serves as an *in vivo* model for condensin-mediated gene regulation.

## Methods and Materials

### Sequence alignment generation

Protein sequences for SMC proteins were found on Wormbase or Uniprot [78,79]. All sequence information is in Supplemental Table 1. The European Molecular Biology Laboratory’s European Bioinformatics Institute (EMBL-EBL) hosts different alignment algorithm tools that are available for use online [80]. One of the available tools is the multiple sequence alignment tool MUSCLE. MUSCLE stands for multiple sequence comparison by log-expectation [81]. All MUSCLE alignments were outputted in the ClustalW parameters. Identity scores are found at the bottom of the alignment. Residues with perfect alignment are given stars, residues with near perfect alignment are given a dot (scoring over 0.5 on the Gonnet PAM 250 Matrix), and residues with highly similar identity are given a colon (scoring under 0.5 on the Gonnet PAM 250 Matrix). Residues with little to no similarity have no indicating mark.

### Cloning and plasmid construction

DPY-27 constructs used in this study: DPY-27 WT N-term: amino acids (aa) 89-282 and DPY-27 WT C-term 1179-1362; DPY-27 EQ substitutes E1275 with Q. MIX-1 construct consisted of N-term: aa 1-176 and C-term: 1038-1229. SMC-4 construct consisted of N-term: aa 91-279 and C-term: aa 1217-1410. Each DNA sequence was found on Wormbase and then codon-optimized for expression in *E. coli*. All primers and synthetic gBlock gene fragments were purchased from IDT and these sequences are listed in Supplemental Table 2. All restriction enzymes were purchased from NEB. The pET-Duet-1 and pET-SUMO plasmid was gifted by JK Nandakumar. DPY-27 N-term (aa 89-282) was cloned into pET-SUMO using BamHI-HF and SalI-HF. The entire fragment of 10xHis-SUMO-DPY-27 (N-term) was then cloned into the multiple cloning site 1 (MCS1) of pET-Duet-1 using NcoI-HF and SalI-HF (which already had DPY-27 WT C-term (aa 1179-1362) in MCS2 using NdeI and XhoI). For EQ DPY-27, the exact same 10xHis-SUMO-DPY-27 (N-term) fragment was cloned into MCS1 of pET-Duet-1 which had the EQ DPY-27 C-term (aa 1179-1362, E1275Q) in MCS2. MIX-1 N-term was cloned into MCS2 of pET-Duet-1 using NdeI and XhoI, and then MIX-1 C-term was cloned into MCS1 using BamHI-HF and SalI-HF. SMC-4 was cloned exactly as DPY-27, with 10xHis-SUMO-SMC-4 N-term in MCS1 and SMC-4 C-term in MCS2 of MCS2 of pET-Duet-1. All cloned plasmids were subjected to Sanger sequencing for validation.

### Protein expression and purification

The SMC ATPase heads (DPY-27 WT, EQ DPY-27, MIX-1, and SMC-4) were expressed in BL21 (DE3) *E. coli* cells after induction with 0.1mM isopropyl β-D-1-thiogalactopyranoside (IPTG) at 16°C for 20 hours. For lysis, pellets were resuspended in lysis buffer (25mM Tris HCl pH 7.5, 500mM NaCl, 10mM 2-mercaptoethanol, 0.1µM EDTA, 1mM PMSF, and 1x protease inhibitor [cOmplete Protease Inhibitor Cocktail, Roche]) and sonicated. The lysate was then clarified via centrifugation at 22,000 x g for 30 min at 4°C. The clarified supernatant was incubated with nickel-NTA agarose beads (Qiagen) for 2 hours at 4°C. Protein bound beads were collected by gravity flow and washed twice with wash buffer (25mM Tris-HCl pH7.5, 150mM NaCl, and 13.6mM 2-mercaptoethanol). Beads were washed then with a high-salt buffer (25mM Tris-HCl, 300M NaCl, 13.6mM 2-mercaptoethanol, and 22.7mM imidazole), before elution (wash buffer supplemented with 300mM imidazole pH 8.0). The 10xHis-SUMO tag was cleaved from DPY-27 WT, DPY-EQ, and SMC-4 by incubating the eluted protein with Ulp1 protease (1% of total eluted protein) for 20 minutes at room temperature. All ATPase heads were further subjected to size exclusion chromatography (Superdex 75 column, GE Healthcare) in buffer containing 25mM Tris-HCl pH 7.5, 100mM NaCl, and 1mM 1,4-dithiothreitol (DTT). Purified ATPase heads were aliquoted in small volumes, flash frozen in liquid nitrogen, and stored at –80° C. MIX-1 and DPY-27 ATPase heads were validated by proteomics analysis, performed by the Proteomics Resource Facility, a part of the Biomedical Research Core Facilities at the University of Michigan.

### ATPase Assay

All reactions were set up with 1mM ATP and 20µM DNA (a purified PCR product of 588 bp amplified with primer sequences listed in Table S2) in a reaction buffer (25mM Tris-HCl pH 7.5, 100mM NaCl, 1mM 1,4-dithiothreitol (DTT), 0.1mg/ml BSA, and 20mM MgCl_2_). 2.5mM of each ATPase head was used for each reaction. Reactions were incubated in a white bottomed 96-well plate for 15 minutes at room temperature, following a 30 second spin at 4000 x g and orbital shake for 2 min on medium (Spectramax iD3, Molecular Devices). Reaction was then quenched and unreacted ATP was depleted using the ADP-Glo reagent from the ADP-Glo kit (Promega) for 40 minutes at room temperature, following a 30 second spin at 4000 x g and orbital shake for 2 min on medium. ADP resulting from the hydrolysis reaction were converted back to ATP, then used to convert luciferin to light using the Kinase Detection Reagent from the ADP-Glo kit for 1 hour at room temperature, following a 30 second spin at 4000 x g and orbital shake for 2 min on medium. Relative luminescence units (RLU) were measured on a GloMax (Promega, 0.5 second integration). Three technical replicates per assay were performed. A one-way ANOVA followed by multiple comparisons was performed to determine statistical significance between conditions. Bar graphs represent the average RLU per 3 replicates with standard deviation.

### *Caenorhabditis elegans* strains and maintenance

Strains were grown under standard conditions as described in [82]. Worm strains are listed in Supplemental Table 3. Worms were fed OP50 unless stated otherwise.

#### Generation of *dpy-27 (cld21)*

We followed the CRISPR/Cas9 protocol generated by the Seydoux lab [83] to introduce the E1275Q mutations in the *dpy-27* gene. All guide RNAs and oligos were obtained from IDT. The E1275Q mutation was generated by substituting G3822 (in cDNA) to a C, changing the GAA codon to CAA in the repair template. To genotype for the E1275Q edits, we PCR amplified small regions centered on the mutants and digested the PCR product with Sau3AI. See Table S2 for all sequence details.

#### Generation of *dpy-27 (cld22*)

We followed the CRISPR/Cas9 protocol generated by the Seydoux lab [83], however the spontaneous NHEJ repair did not include the supplied repair template. Instead, repair resulted in insertion and premature stop codon in exon 11 of *dpy-27*. To genotype for cld22 Indel, we PCR amplified small regions centered around the insertion. Insertion is 57bp longer than wildtype. See Table S2 for all sequence details.

#### Generation of FLAG::*dpy-21*

We followed the CRISPR/Cas9 protocol generated by the Seydoux lab [83] to introduce a 3xFLAG tag after the fifth amino acid of the *dpy-21* coding sequence. All guide RNAs and oligos were obtained from IDT. To genotype for the 3xFLAG insertion, we PCR amplified the region surrounding the insert. The PCR product from the edited allele is 84 bp longer than from the wild type allele. See Table S2 for all sequence details.

### Brood Counts

For brood counts, L4 hermaphrodites of the appropriate strains were picked onto 30mm plates. Hermaphrodites were moved daily, and the number of total progeny (eggs and hatched larvae) was counted after the parent was moved. The number of dead eggs was counted after 24 hours, and the number of L4 (or older) progeny was counted on the 3^rd^ day from original parent being set down on plate. Counts were done until parental worm reached end of reproductive span (counted as two consecutive days with 0 progeny laid) or when parent worm died. All parental worms were genotyped for confirmation. Statistical significance was determined by a one-way ANOVA.

### Zygotic Rescue Assay

For the zygotic rescue assays, 5 L4 hermaphrodites of the appropriate strains were picked onto 30mm plates (one L4 hermaphrodite per plate) along with 3-4 *him-8 mIs10* males per plate. 5 additional L4 hermaphrodites of the appropriate strains were picked onto 30mm plates alone for control. Hermaphrodites were allowed to lay eggs for 48 hours and then removed for PCR genotyping. Males were also removed at the same time. Total progeny laid on plates were counted at this time. The number of dead eggs were counted after 24 hours. 72 hours after parental removal, all progeny that reached L4 and later were counted, as well as number of L4 progeny, hermaphrodite vs male, and GFP+ (cross progeny) vs non-GFP+ (self-progeny). For crosses, the few number of non-GFP+ self progeny was subtracted from the total so only cross progeny are shown on the graphs. At least 3 L4 progeny were selected for PCR genotyping. Larval viability was scored based on number of progeny that reached L4 or later at the 72 hours post parental removal divided by total progeny laid at parental removal. Statistical significance was determined by student’s t-test between control hermaphrodite broods and crossed hermaphrodite broods. A minimum of two trials were included in the results, with a final n of 10 parent hermaphrodites per condition.

### Immunofluorescence

Immunofluorescence (IF) experiments were performed as described in [16]. One day post-L4 worms were dissected in 1x Sperm Salts (50 mM Pipes pH 7, 25 mM KCl, 1 mM MgSO4, 45 mM NaCl and 2 mM CaCl2) and fixed in 2% formaldehyde for 5 min with a coverslip in a humidity chamber filled with PBST (1x PBS with 0.5 % triton X-100). Slides were then frozen on dry ice for at least 15 minutes. Coverslips were removed and slides were then washed in PBST three times for 10 minutes each. Slides were subsequently incubated with primary antibodies for overnight at room temperature. Slides were then washed in PBST (3 x 10 min each), before incubation in secondary antibody for 1 hour at 37°C. Slides were washed in PBST (2 x 10 min each), followed by a 10-minute incubation of DAPI diluted in PBST. Slides were mounted with Vectashield (Vector Labs).

For detergent extraction experiments, one day post-L4 worms were dissected in 1x Sperm Salts with 1% triton (50 mM Pipes pH 7, 25 mM KCl, 1 mM MgSO4, 45 mM NaCl and 2 mM CaCl2) and incubated in humidity chamber for up to 3 minutes. Then IF protocol from above was followed, starting with fixation with 2% formaldehyde.

### Antibodies Used

Primary antibodies used include Rabbit anti-DPY-27 at 1:100 [16], Goat anti-CAPG-1 at 1:250) [84], Rabbit anti-H4K20me1 at 1:500 (Abcam ab9051), Rabbit anti-H4K20me2 (Abcam 9052), Mouse anti-FLAG at 1:1000 (Millipore Sigma F1804), and Rat anti-HTZ-1 at 1:100 [85].

Secondary antibodies used include Donkey anti-rabbit-FITC at 1:100, Donkey anti-Mouse-FITC at 1:100, Donkey anti-Rabbit-Cy3 at 1:100, Donkey anti-Goat-Cy3 at 1:100, and Donkey anti-rat-Cy3 at 1:100 from Jackson Immunochemicals.

### Chromosome PAINT probe synthesis

Yeast artificial chromosomes (YAC) containing sequences that corresponded to either the X chromosome or Chromosome I were amplified via degenerate primer PCR as described in [84,86]. Fluorescent dCTP-Cy3 (GE) was incorporated along with standard dNTPs with random priming (Prime-a-Gene Kit, Promega). Probes were precipitated in 95% ethanol, salmon sperm DNA (Roche), and 3M sodium acetate overnight at -20°C. After washing with 70% ethanol and drying for at least 30 minutes at room temperature, each probe was resuspended in either Hybrisol VII (MP Biomedicals) or 80ul hybridization buffer as described in [87]. Briefly, 80µl hybridization buffer consists of 5.76µl TEN buffer (1M Tris-HCl pH7.5, 0.5M EDTA, 1M NaCl), 40µl 100% deionized formamide, 8µl 10% blocking reagent (10% w/v solution in 1x maleic acid buffer), 16µl 50% dextran sulfate, and nuclease free water up to 80µl.

### Fluorescent *in situ* Hybridization (FISH)

24 hours post L4 worms were dissected in 1x Sperm Salts (50 mM Pipes pH 7, 25 mM KCl, 1 mM MgSO4, 45 mM NaCl and 2 mM CaCl2) and fixed in 2% formaldehyde for 5 min with a coverslip in a humidity chamber filled with 2xSSC (30mM sodium citrate, 300mM NaCl), 50% formamide (Roche). Slides were then frozen on dry ice and kept frozen for at least 20 minutes. Coverslips were removed and slides were then washed in PBST three times for 10 minutes each. Slides were then dehydrated in an ethanol series: 70%, 80%, 95% and 100% for 2 minutes each. After allowing slides to dry for at least 10 minutes or until all alcohol had evaporated, 10µl of Cy3-labelled probe was applied with a coverslip and heated to 95°C for 3 minutes. Identical probe aliquots were used on wildtype and mutant strains for each experiment. Slides were then incubated overnight at 37°C in a warm humid chamber (filled with 2xSSC, 50% formamide). The following day, slides were subjected to a series of washes: 3 washes of 2xSSC, 50% formamide for 5 minutes each at 39°C, 3 washes of 2xSSC for 5 minutes each at 39°C, and 1 wash of 1xSSC for 10 minutes at 39°C. The final step was a 10-minute incubation of 4xSSC with DAPI at room temperature. Slides were then mounted with Vectashield (Vector Labs).

### Combined FISH and IF

One day post-L4 hermaphrodites were processed for FISH as above. After the 1xSSC wash, slides were washed once in PBST for 10 minutes, and then IF protocol was performed, beginning with incubation of primary antibody.

### Imaging

Images were collected with a Hamamatsu ORCA-ER digital camera mounted on an Olympus BX61 epi-fluorescence microscope with a motorized Z drive. All images were taken with the 60x APO oil immersion objective. Z-stack series were collected at 0.2 um increments. Gut nuclei images shown are maximum intensity projection images summed from ∼8 um. Embryonic nuclei images shown are maximum intensity projection images summed from ∼0.6 um. All images shown were created by exporting the 16-bit TIFF z-stack series and using ImageJ (Fiji) to create projections. Projection images were then adjusted for brightness and contrast in Photoshop (Adobe), with all adjustments consistent between genotypes. Final overlays were created in ImageJ (Fiji).

### Nuclear Volume Quantification

Slidebook 5 (Intelligent Imaging Innovations) was utilized for volumetric quantification. Using the segment mask functions, individual masks for DAPI (DNA) and Cy3 (FISH) were drawn on each nucleus. Using the mask statistics function with DAPI as the primary mask and Cy3 as the secondary mask, the morphometrix (voxels) and crossmask (voxels) values were generated. Chromosome occupancy/volume was calculated by dividing the crossmark value by the morphometrix value (corresponding to Cy3/DAPI). A minimum of 20 nuclei per genotype were quantified for each experiment. These nuclei were collected from an average of 6 worms per genotype, with an average of 4 nuclei per worm. A one-way ANOVA with multiple comparisons was conducted in Prism (GraphPad) to compare statistical significance between genotypes.

### Immunofluorescence and FISH Overlap Quantification

Slidebook 5 (Intelligent Imaging Innovations) was utilized for overlap quantification. Using the segment mask functions, individual masks for DAPI (DNA), FITC (Immunofluorescence) and Cy3 (FISH) were drawn on each nucleus. Using the mask options menu, a combined mask with voxels present in both FITC and Cy3 masks was generated. Using the mask statistics function with DAPI as the primary mask to isolate signal to within the nucleus only and either the combined mask or FITC as the secondary mask, the crossmask (voxels) values were generated. Overlap was calculated by dividing the combined crossmask value by the FITC crossmask value (corresponding to overlap/total FITC signal). A minimum of 20 nuclei per genotype were quantified for each experiment. These nuclei were collected from an average of 7 worms per genotype, with an average of 3 nuclei per worm. A one-way ANOVA with multiple comparisons was conducted in Prism (GraphPad) to compare statistical significance between genotypes.

## Results

### DPY-27 is capable of hydrolyzing ATP

Using protein sequences from *C. elegans*, human, xenopus, and other species (details in Supplemental Table 1), we used MUSCLE [80,81] to generate sequence homology alignments to understand how DPY-27 compares to other SMC proteins. From the resulting alignments (Supplemental Figure 1), we concluded that DPY-27 has all the required domains that make up the SMC protein ATPase head. Therefore, we tested whether this head is functional by subcloning the ATPase domains of DPY-27 and expressing them in *Escherichia coli* BL21 cells for purification. We also subcloned, expressed, and purified the ATPase domains of MIX-1, which is the heterodimeric partner of DPY-27 in condensin I^DC^ and a mutant version of DPY-27, where we mutated the essential glutamate in Walker B to a glutamine (EQ) that has been shown in numerous organisms to abolish ATPase function [15,26,39,47,48]. We also expressed and purified the ATPase domains of SMC-4 as a comparison. Using an *in vitro* ATPase assay, we found that the combination DPY-27 and MIX-1 is capable of hydrolyzing ATP and at a rate comparable to the mitotic pair of MIX-1 and SMC-4 (Figure 1D). Moreover, the EQ mutation of DPY-27 does abolish ATPase function. We also tested each ATPase head alone (Supplemental Figure 2). Interestingly, both SMC-4 and DPY-27 are capable of hydrolyzing ATP alone. In contrast, the EQ mutant of DPY-27 was barely functional. Therefore, it can be assumed that DPY-27 is a functional ATPase and the function of the essential glutamate in the Walker B domain is conserved.

### The ATPase function of DPY-27 is required for C. elegans viability

While the *in vitro* biochemical data demonstrates that DPY-27 is a functional ATPase, it does not answer whether that function is required for DC. Using CRISPR-Cas9, we generated two *dpy-27* mutants: *dpy-27 EQ*, where we mutated the key glutamate (E) to glutamine (Q) in the endogenous locus; and *dpy-27 Indel,* which results in a premature stop codon and truncated protein. Both mutants were generated in the same experiment, but the *dpy-27 Indel* mutants resulted from a lack of homologous repair. For both mutations, mutants worms appear phenotypically similar. Heterozygous worms are phenotypically similar to wildtype (Figure 2, top left). These worms have inherited maternally contributed wild type DPY-27 protein as well as RNA and are able to produce wild type DPY-27 protein, and thus are referred to as m+z+ (maternal +, zygotic +). Homozygous mutant worms descended from heterozygous parents have inherited wild type protein and RNA from their mothers but are unable to produce wild type protein themselves (m+z-) [58,61,88]. The maternal contribution allows the m+z- worms to survive to adulthood, but they develop Dpy (dumpy, short and fat) (Figure 2, top middle). Most of the progeny of these mutants (lacking both maternal and zygotic contribution of DPY-27, m-z-) arrested as larvae (Figure 2, top right). These phenotypes are consistent with previous findings on *dpy-27* and other DCC mutants [56,58,60–62,65,89].

**Figure 2.**
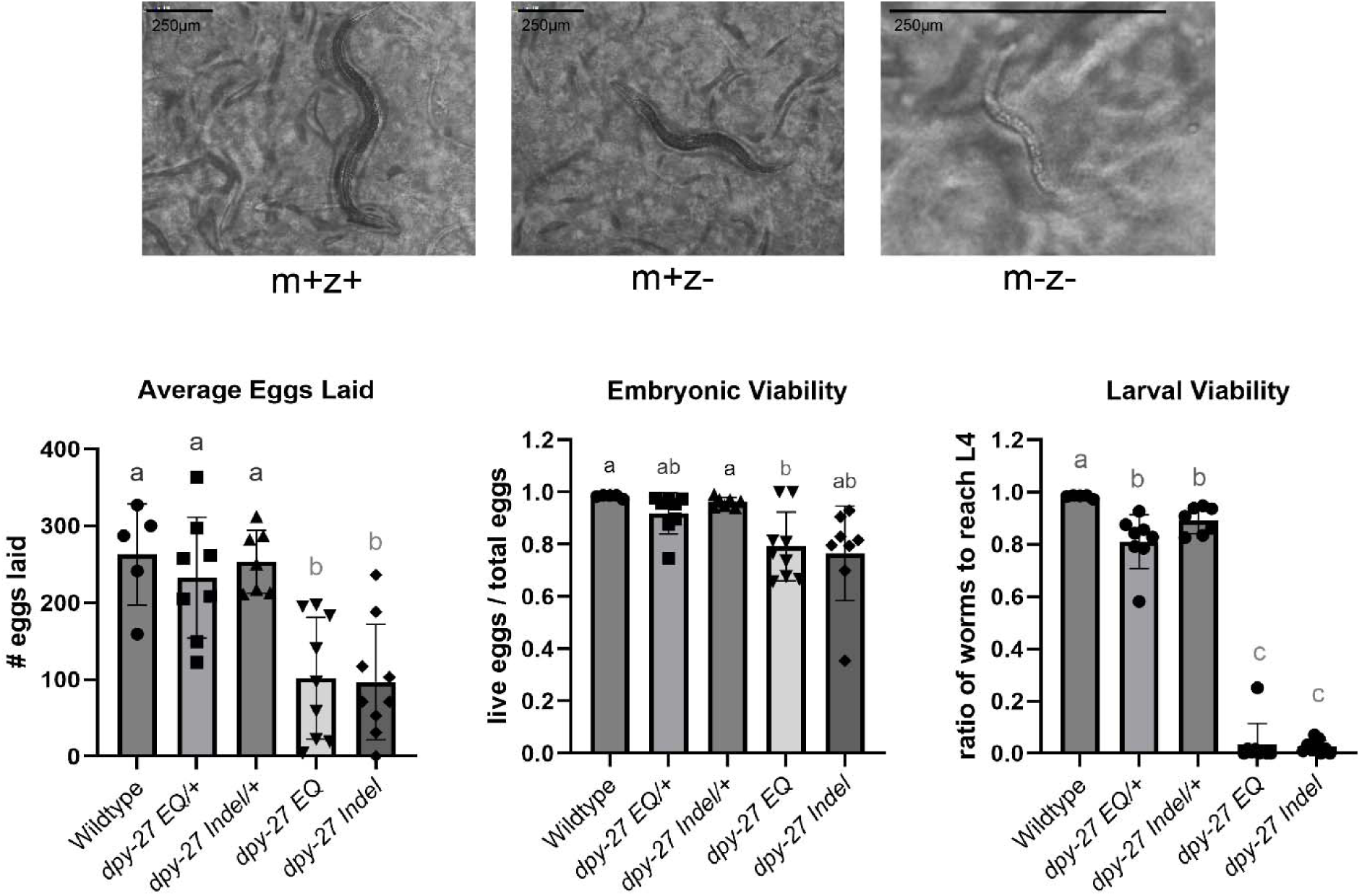
*dpy-27* mutant worm phenotypes. **(Top)** Representative images of age-matched worms at 120 hours post hatching. Phenotypes of the mutants are similar; *dpy-27 EQ* m+z- and m-z- mutants are shown. Scale bars are 250 µm. **(Bottom)** Average eggs laid, embryonic viability, and larval viability of progeny for *dpy-27* mutants. Embryonic viability was determined by “number of live eggs/total eggs laid”. Larval viability was determined by “number of worms to reach L4/total eggs laid”. “n” number for each genotype follows: wildtype = 5, *dpy-27 EQ* = 9, *dpy-27 Indel* = 9, *dpy-27 EQ/+* = 8, and *dpy-27 Indel/+* = 7. Different letters indicate statistically significant differences determined through a Brown-Forsythe ANOVA and Dunnett’s T3 multiple comparisons. Any repeated letter demonstrates no significance between groups. See Table 5 for all p-values.

To evaluate how these mutations effect fitness, we quantified the number of eggs laid, the percentage of embryos that hatched into larvae (embryonic viability), and the percentage of progeny that reached L4 or adults stages (larval viability) (Figure 2, bottom). We compared the eggs laid and the progeny produced by wildtype hermaphrodites, heterozygous hermaphrodites for both *dpy-27* mutations, and first-generation m+z- homozygous hermaphrodites for both *dpy-27* mutations. Wildtype worms and heterozygous worms for both *dpy-27* mutants laid a comparable number of eggs, averaging near 250. In contrast, both *dpy-27 EQ* and *dpy-27 Indel* homozygous m+z- worms laid significantly fewer, averaging about 100 eggs, indicating that *dpy-27* mutants have reduced fertility. Embryonic viability for both homozygous *dpy-27 EQ* and *dpy-27 Indel* worms was lower than the wildtype and heterozygous worms, but most of the progeny hatched for all genotypes. The difference in embryonic viability was not statistically significant for *dpy-27 Indel* worms compared to other genotypes. One reason for lack of statistical significance was that the *dpy-27 Indel* worms had a very broad range of embryonic viability, with one worm being sterile, another had less than 40% viability with most of its progeny dead before hatching, and the rest had over 70% of the laid eggs hatched. The differences in larval viability was more significant. Very few of the m-z- progeny of both *dpy-27 EQ* and *dpy-27 Indel* homozygous m+z- worms developed into L4 or adult hermaphrodites. Most of these animals died or arrested as L1-L2 larvae, indicating that these mutations result in a maternal effect lethal phenotype. Interestingly, both *dpy-27 EQ/+* and *dpy-27 Indel/+* heterozygous worms had decreased larval viability compared to wildtype, and this is most likely due to the 25% homozygous mutant progeny they produce, which are slower developing and did not reach the L4 stage in the time frame of the experiment. However, based on the result that about 80% of the progeny of heterozygous worms reached this stage, we can conclude that heterozygous worms have no viability or fertility defects. These results suggest that neither *dpy-27* mutation is acting like a dominant negative.

The decreased embryonic viability and increased larval arrest in the m-z- progeny of both *dpy-27 EQ* and *dpy-27 Indel* homozygous worms was not unexpected and is consistent with reported phenotypes of other severe loss of function *dpy-27* mutants [56]. While the m+z- generation survives due to maternally deposited wild type RNA and/or protein of DCC members into the egg, the m-z- offspring die or arrest as larvae because they do not receive wildtype protein or RNA [58,61,88]. However, there is a possibility that transcription of wildtype *dpy-27* by the embryo could rescue the lack of maternally loaded wildtype DPY-27 protein. To test this, we used a zygotic rescue assay where a wildtype copy of *dpy-27* was paternally donated. To do so, we set up each homozygous mutant m+z- hermaphrodites either alone or with three males homozygous wild type for *dpy-27* and bearing a GFP+ marker (homozygous for the mIs10 transgene) (Figure 3A). Any progeny that expressed GFP were cross-progeny and considered to have a wildtype copy of DPY-27, making them m-z+. Self-progeny (lacking GFP expression) were rare, but were counted separately from the cross-progeny and not included in the graphs.

**Figure 3.**
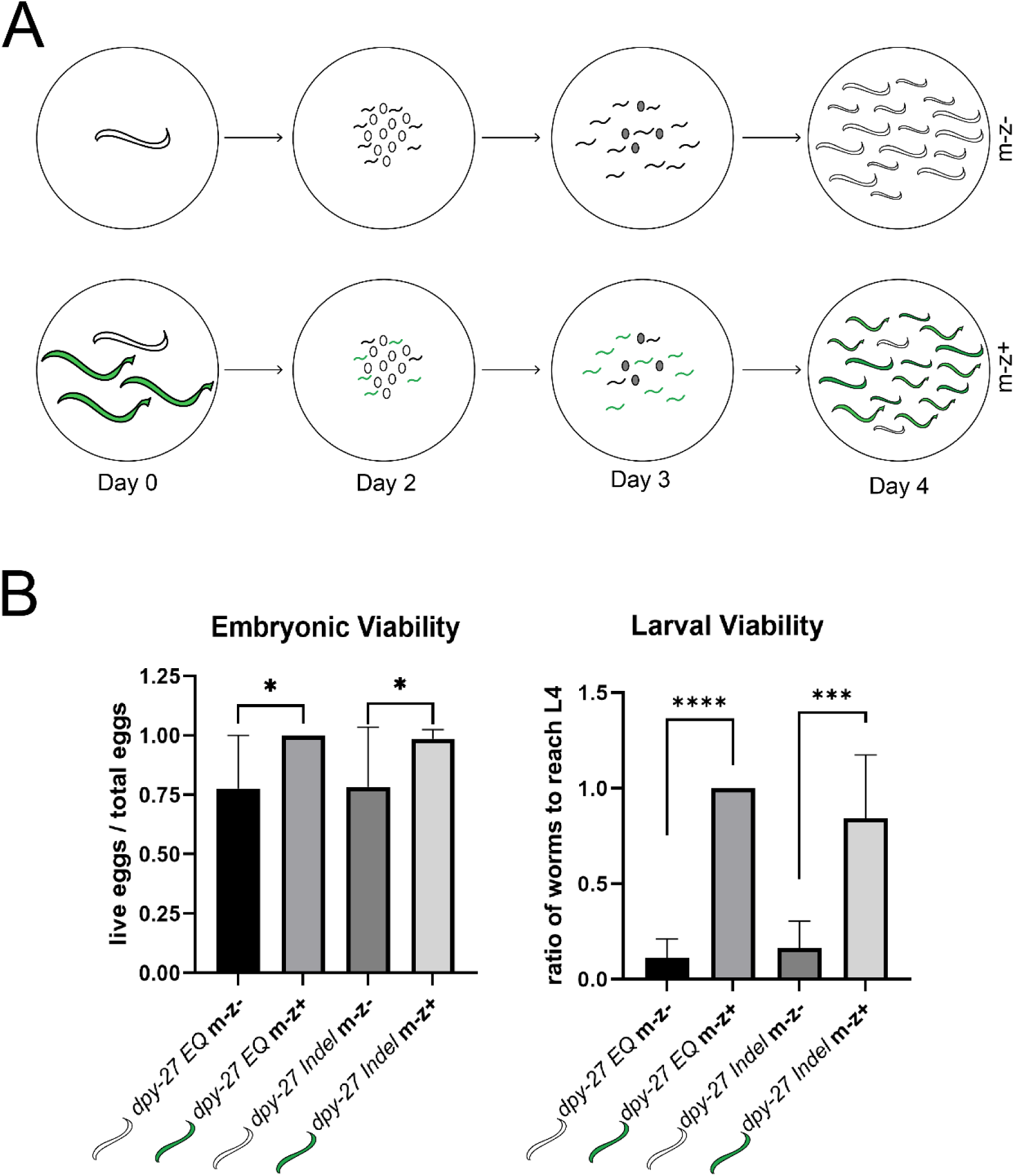
Zygotic Rescue Assay. **(A)** Zygotic rescue assay schematic shows experimental set-up. A single mutant m+z- L4 hermaphrodite was allowed to either self-fertilize to produce “m-z-” offspring (first row) or paired with 3 males to produce “m-z+” offspring (second row) for 2 days. On day 2, parents were removed for genotyping and the number of progeny laid was counted. The number of dead eggs was counted on Day 3 and the number of progeny reaching L4 was counted on day 4. **(B)** Embryonic and larval viability for *dpy-27* “m-z-“ and *dpy-27* “m-z+” worms. Embryonic viability was determined by “number of live eggs/total eggs laid”. Larval viability was determined by “number of worms to reach L4/total eggs laid”. Number of parent hermaphrodite used per genotype follows: *dpy-27 EQ m-z-* = 6, *dpy-27 EQ m-z+* = 3, *dpy-27 Indel m-z-* = 8, and *dpy-27 Indel m-z+* = 7. Statistical significance for embryonic viability was determined by a two-tailed Mann Whitney test (*EQ m-z-* vs *EQ m-z+* p = 0.0238; *Indel m-z-* vs *Indel m-z+* p = 0.0263). Statistical significance for larval viability was determined by an unpaired student’s t-test (*EQ m-z-* vs *EQ m-z+* p < 0.0001, *Indel m-z-* vs *Indel m-z+* p = 0.0001).

Both the *dpy-27 EQ m-z+* and *dpy-27 Indel m-z+* progeny had 100% embryonic viability and nearly 100% of the progeny reached the final larval stage L4 (Figure 3B). In the crosses, about half (47-50%) of the *m-z+* progeny were male as expected. The rescued *m-z+* hermaphrodites were healthy and fertile. This demonstrated that zygotic transcription of *dpy-27* is indeed sufficient to rescue lack of maternally loaded protein. Moreover, it also demonstrates that zygotic transcription of *dpy-27* begins early enough in development to prevent embryonic arrest and that a single copy of wildtype *dpy-27* is sufficient for this rescue. Taking both the fitness and zygotic rescue results, our genetic analysis demonstrated that a single point mutation that just abolishes the ATPase activity of *dpy-27* results in mutant phenotypes comparable to previously characterized null mutants that lost the entire gene [56].

### ATPase-Defective DPY-27 has varied DNA-binding ability

To understand how the EQ mutation affects DPY-27 function at the molecular level, we first assayed whether mutant protein is produced. In a study that produced an EQ mutant DPY-27 protein from a heat shock inducible transgene in a genetic background that had a wildtype copy of the *dpy-27* gene, the GFP-tagged EQ-DPY-27 protein degraded over time and did not colocalize with other members of the DCC [77]. In our homozygous mutant worms, there is no wildtype protein produced. Thus, we were able to investigate how the mutant protein behaves when no wildtype protein is present. Using immunofluorescence (IF), we stained for DPY-27 and CAPG-1, another member of the DCC (Figure 4A). In wildtype gut nuclei, DPY-27 and CAPG-1 colocalize and their localization is limited to a portion of nucleus, as has been shown before for members of condensin I^DC^ bound to the X chromosomes [16,59,61,69]. *dpy-27 Indel* nuclei were completely different, with no visible DPY-27 staining and very little CAPG-1, phenocopying a published DCC mutant, *dpy-28 null*. This phenotype has been documented before for other null (or severe loss of function) mutations in genes encoding condensin I^DC^ subunits. In the absence of one subunit, the complex is not stable [59,60,62,68,69,90]. Therefore, we assume that in *dpy-27 Indel* mutants, the truncated protein is not accumulated at a level high enough to be detectable by IF, leading to loss of X localization for other complex members as well. Moreover, the lack of staining in the *dpy-27 Indel* mutants demonstrates that the maternally loaded wildtype protein is not detectable by the adult stages. Interestingly, in *dpy-27 EQ* mutants, all of the nuclei showed detectable DPY-27 protein, which colocalized with CAPG-1. A majority of these nuclei had staining over all of the nucleus (Figure 4A, row 2). A smaller set of nuclei, in contrast, had staining that was more restricted and reminiscent of X chromosome localization (Figure 4A, row 3). These results suggest that the EQ mutant protein is more stable than the indel, and it may be able to be incorporated into the condensin I^DC^ complex.

**Figure 4.**
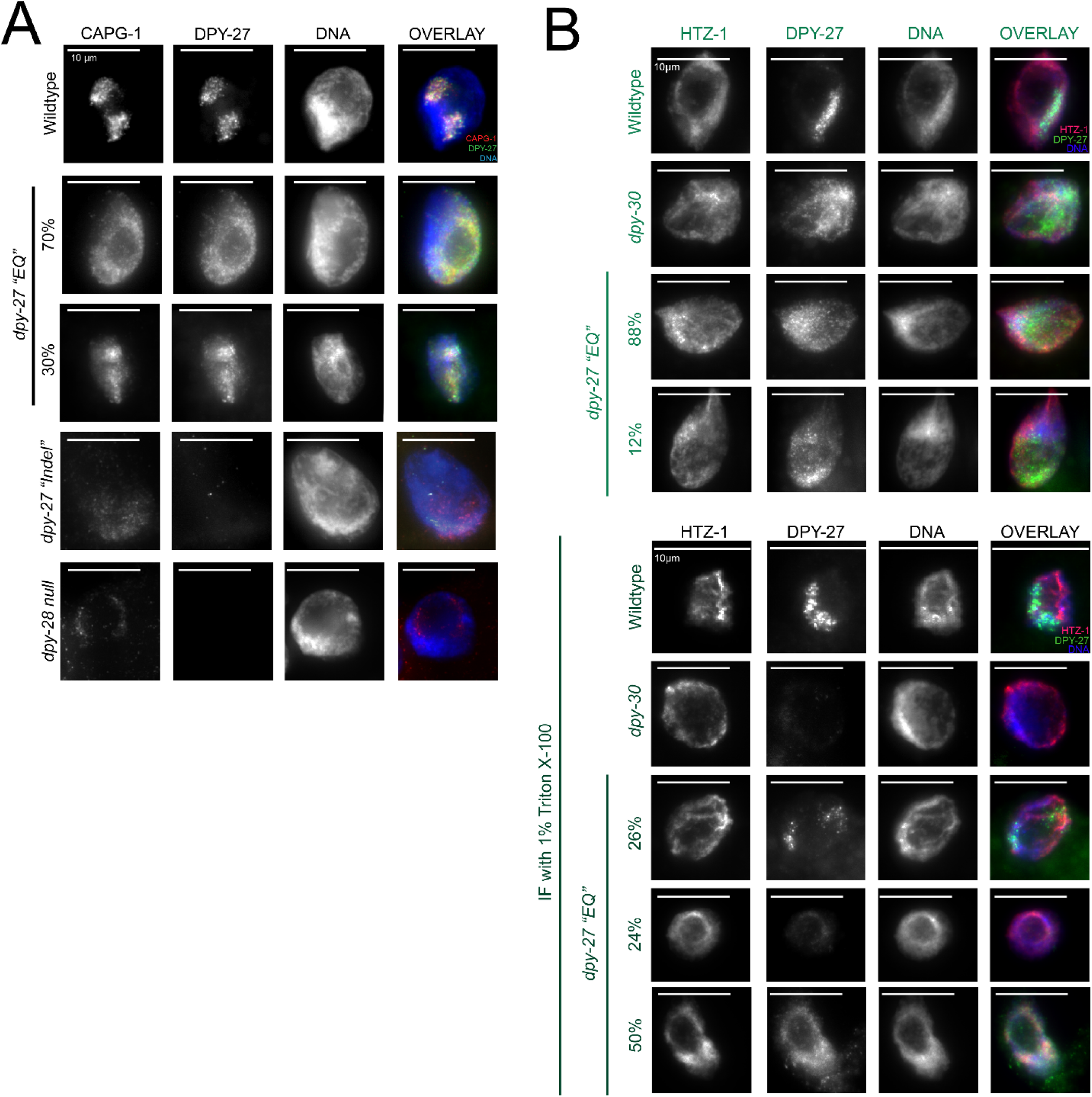
DPY-27 localization and DNA binding in differentiated gut nuclei. **(A)** Intestinal nuclei of young adult hermaphrodites of each genotype were stained with antibodies against DPY-27 and CAPG-1. In wildtype, DPY-27 is visible and colocalizes with CAPG-1 on X chromosomes (top row). In *dpy-27 “EQ”* mutant nuclei, DPY-27 is visible and colocalizes with CAPG-1. In the majority of nuclei staining is visible all over the nucleus (second row, n = 17/25), while a smaller percentage of nuclei have a less diffuse staining pattern, indicating some degree of subnuclear localization (third row, n = 8/25). In contrast, *dpy-27 “Indel”* and *dpy-28 null* nuclei have no staining for DPY-27 and very weak CAPG-1 (fourth and fifth rows). **(B)** Intestinal nuclei were stained with antibodies against DPY-27 and HTZ-1 (staining control) after dissection in the presence or absence of a detergent. In wildtype, both DPY-27 and HTZ-1 remain DNA-bound, whether in the presence of the detergent or not (row 1 vs row 5). *dpy-30* has DPY-27 protein visible all over the nucleus in absence of detergent (row 2), but no staining in the presence of detergent (row 6). In *dpy-27 “EQ”* mutants in the absence of detergent, there are two phenotypes (rows 3-4), with protein all over the nucleus, or some degree of subnuclear localization. In the presence of detergent, these phenotypes are even more apparent, with some nuclei having completely lost signal, others maintaining overall or restricted staining (rows 7-9, n = 9/34, n = 8/34, n = 17/34 respectively).

The all-over staining of DPY-27 is reminiscent of condensin I^DC^ staining patterns in *dpy-30* mutants [59,61,68,69]. In *dpy-30* mutants, the DCC is present but unable to strongly bind to the X chromosomes, leading to diffuse nuclear staining [68]. We previously showed that detergent extraction can be used to remove DCC proteins that are weakly DNA bound, leaving strongly DNA-bound proteins unaffected [68]. To test whether DNA binding is compromised in the *dpy-27 EQ* mutants, we performed IF again, but with an additional detergent extraction step. This time, we stained for DPY-27 and HTZ-1, which is a histone 2A variant and therefore strongly DNA-bound. Without detergent, wild type nuclei show restricted DPY-27 staining, while in *dpy-30* mutants DPY-27 staining is visible all over the nucleus. In the *dpy-27* EQ mutant, we again see the two phenotypes of staining previously observed (Figure 4B, rows 3-4). When 1% triton is included in the dissection buffer, we found that HTZ-1 is unchanged in all genotypes, as expected. DPY-27 is strongly bound in wildtype nuclei but completely lost in *dpy-30* nuclei, consistent with previous findings [68]. In contrast, there is a range of phenotypes in *dpy-27 EQ* nuclei. Some nuclei lose all staining (Figure 4B, row 8), some nuclei maintain all-over staining of DPY-27 (row 9), and some nuclei have staining restricted, resembling wildtype DPY-27 bound to the X (row 7). This wide distribution was not an artifact of uneven detergent penetration as many of the different phenotypes were found within the same worm. Therefore, we conclude that without ATPase function, there are two populations of DPY-27; one population is strongly DNA bound while the other is not.

### Loss of DPY-27 ATPase function results in X-chromosome decompaction

We next assessed how the Indel and EQ mutations in DPY-27 impact X-chromosome condensation. X-chromosome condensation, measured as the volume occupied by the X chromosomes in the nucleus (X-chromosome volume/nuclear volume), is a feature of dosage compensated X chromosomes that is attributed to the activity of condensin I^DC^ [91]. Using whole chromosome PAINT fluorescent *in situ* hybridization (FISH), we assayed condensation of the X chromosomes in gut nuclei (Figure 5A). In wildtype worms, the X chromosomes are tightly compacted and located at the nuclear periphery, as seen before [84,91], Both *dpy-27 EQ* and *dpy-27 Indel* mutants have decondensed X chromosomes that occupy more volume in the nucleus, and the degree of decondensation was equally severe in the *dpy-27 EQ* mutants as it was in the *dpy-27 Indel* mutants (Figure 5B). As a control, we also checked whether autosomes were affected by the mutations, and we assayed condensation of chromosome I (Figure 5C). As we hypothesized, chromosome I is unaffected by either mutation in *dpy-27* (Figure 5D), as DPY-27 function is restricted to the X chromosomes [59,74,92]. Our results suggest that the loss of ATPase function in DPY-27 leads to as severe defects in X chromosome condensation as a truncation mutation of DPY-27 that does not make detectable protein.

**Figure 5.**
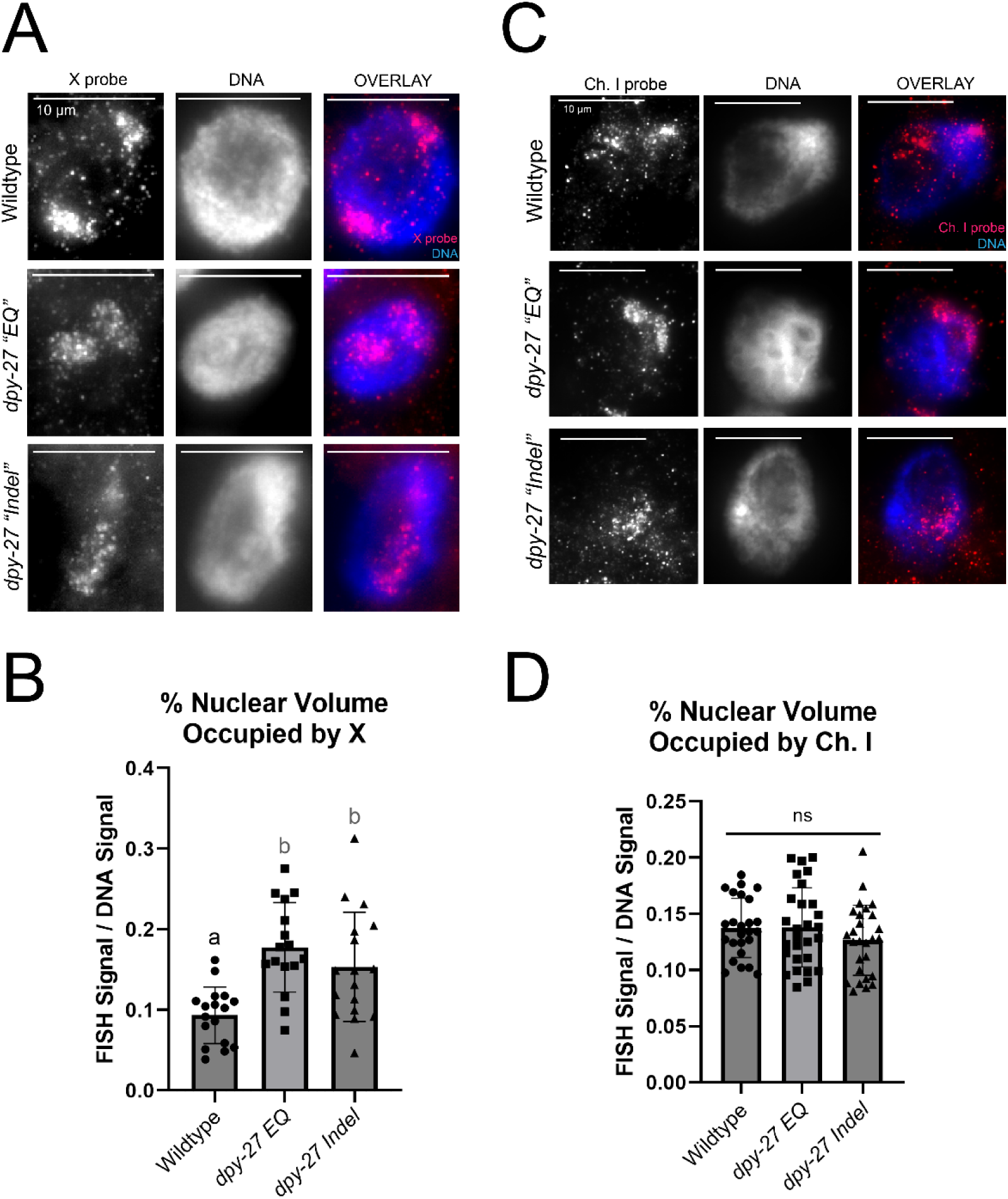
Chromosome compaction in *dpy-27* mutants. **(A)** Using Whole X-PAINT FISH, the X chromosomes in differentiated gut nuclei of age-matched hermaphrodites were assayed for compaction. In wildtype gut nuclei, the X chromosomes are tightly condensed (top row). In both *dpy-27 “EQ”* and *dpy-27 “Indel”* mutant nuclei, the X chromosomes expand significantly in volume (middle and bottom). **(B)** Quantification of % nuclear volume occupied by X chromosomes. Error bars indicate standard deviation. Statistical significance was determined by a one-way ANOVA followed by Šídák’s multiple comparisons test. P-values are: N2 vs *dpy-EQ* p = 0.002, N2 vs *dpy-27 Indel* p = 0.0073, *dpy-27 EQ* vs *dpy-27 Indel* p = 0.4986. **(C)** Using Whole PAINT FISH, chromosome I in differentiated gut nuclei of age-matched hermaphrodites were assayed for compaction. Chromosome I is similarly compacted in wildtype, *dpy-27 “EQ”*, and *dpy-27 “Indel”* gut nuclei. **(D)** Quantification for % nuclear volume occupied by chromosome I, normalized to nuclear size. Error bars indicate standard deviation. Statistical significance was determined by a one-way ANOVA followed by Šídák’s multiple comparisons test. P-values are: N2 vs *dpy-EQ* p = 0.9999, N2 vs *dpy-27 Indel* p = 0.5084, *dpy-27 EQ* vs *dpy-27 Indel* p = 0.4448.

In the *dpy-27 EQ* mutants, some portion of the DPY-27 is DNA bound (Figure 4B). Therefore, we investigated whether the DPY-27 protein binds to the X chromosomes in *dpy-27 EQ* mutant gut nuclei. We combined the whole chromosome PAINT FISH and immunofluorescence to assay DPY-27 localization (Figure 6). In wildtype gut nuclei, DPY-27 colocalizes clearly with the X chromosomes. Interestingly, DPY-27 in *dpy-27 EQ* gut nuclei is bound to the X chromosomes, but also found elsewhere in the nucleus (Figure 6A, row 2). When quantified, we found that, on average, less than half of the DPY-27 signal was on the X chromosomes in *dpy-27 EQ* gut nuclei, compared to over 75% in wildtype (Figure 6B). We observed a range of phenotypes in *dpy-27 EQ* nuclei and the full range is shown in Supplemental Figure 3. We also attempted to see if there was a correlation between DPY-27 signal on X and the volume of the X chromosomes. We found that in *dpy-27 EQ* mutants, the amount of DPY-27 on the X chromosomes did not correlate with the level of X-chromosome condensation (Figure 6C). Therefore, despite producing DPY-27 protein that can bind the X chromosomes, it is clear that the loss of ATPase function renders the protein to be non-functional.

**Figure 6.**
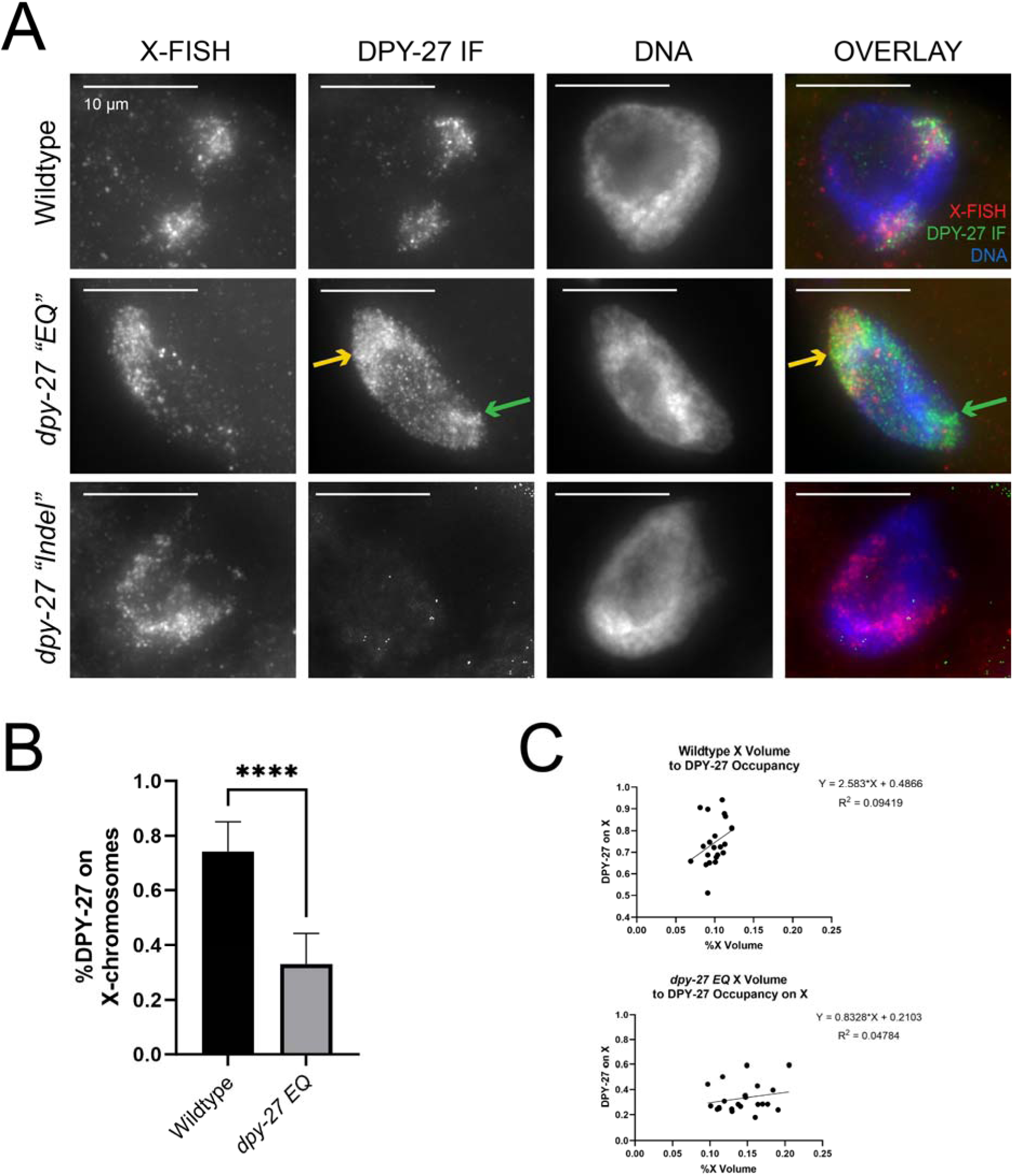
DPY-27 localization on X chromosomes. **(A)** Using Whole X-PAINT FISH and antibodies against DPY-27, localization of DPY-27 in differentiated gut nuclei of age-matched hermaphrodite worms was assayed. In wildtype nuclei, DPY-27 is specifically found on the X chromosomes (top row). In *dpy-27 “EQ”* nuclei, there are two populations of DPY-27, one that is localized to the X chromosomes, and another that is not (middle row, yellow vs green arrows). No DPY-27 protein is visible in *dpy-27 “Indel”* nuclei (bottom row). **(B)** Quantification of the proportion of DPY-27 signal localized to X chromosomes. Error bars indicate standard deviation. Statistical significance was determined by an unpaired student’s t-test with p < 0.0001. **(C)** Correlation of X-chromosome volume and percentage of DPY-27 on X chromosomes was plotted. Best fit linear correlation line and equation with r^2^ are shown.

As a control, we also performed this assay in *dpy-27 EQ/+* heterozygotes (Supplementary Figure 4). While *dpy-27 indel/+* heterozygotes resemble wildtype worms with DPY-27 protein only on the X chromosomes (Supplementary Figure 4A, rows 1 and 4), *dpy-27 EQ/+* heterozygotes had two phenotypes. The first phenotype resembled wildtype with DPY-27 mostly on the X chromosomes, but the second phenotype resembled the homozygous mutants with only a percentage of DPY-27 protein bound to the X chromosomes (Supplementary Figure 4A and B). These worms make both wild type and EQ mutant protein, which is reflected in the DPY-27 localization patterns. We then assayed the nuclear occupancy of the X chromosome and found that there was no difference between the two *dpy-27 EQ/+* heterozygous populations and that their X chromosomes had no decondensation phenotype (Supplementary Figure 4C). From this data, we can conclude that having wildtype DPY-27 is enough for proper X-chromosome condensation and that the presence of the DPY-27 EQ protein does interfere with the wildtype protein’s function. This conclusion is consistent with our genetic analysis above which suggested that the *dpy-27 EQ* mutation does not act like a dominant negative.

### The ATPase-defective DPY-27 cannot colocalize with SDC-2

The localization of the DCC to the X chromosomes is dependent on rex sites. *rex* sites, or *Recruitment element on X* sites, are repeated sequences found on the X chromosomes that directly recruit the DCC to the X chromosome [92–95]. The non-condensin DCC subunit SDC-2 first binds *rex* sites [94] and recruits the rest of the DCC to the X chromosomes [70]. As measured by ChIP-seq, SDC-2 and DPY-27 both have the highest level of occupancy at these sites. To test whether the EQ mutant DPY-27 protein is able to co-localize with SDC-2, we generated a strain that has the *dpy-27 EQ* mutation as well as an SDC-2::FLAG transgene and a mutation at the endogenous *sdc-2* locus (*y74* allele). In this strain, the transgenic SDC-2::FLAG is the only functional SDC-2 protein. The SDC-2::FLAG transgene is functional, as it is able to rescue the hermaphrodite-specific lethality caused by the *sdc-2(y74)* mutation. Using X-chromosome PAINT FISH and antibodies against FLAG, we showed that SDC-2 is localized to X chromosomes in the *dpy-27 EQ* mutants (Supplemental Figure 5). Since SDC-2 is the only DCC subunit able to bind the X chromosomes even when other DCC members are mutated [70], this result was expected.

We then used antibodies against FLAG and DPY-27 to test the degree of colocalization between SDC-2 and DPY-27 (Figure 7A). In SDC-2::FLAG worms which are phenotypically wildtype, FLAG and DPY-27 colocalize. SDC-2 and DPY-27 staining both appear as a cloud of puncta, which are restricted to the X chromosomes. In SDC-2::FLAG worms these puncta often, although not always, colocalize. In the merged image, the overlap is visible as yellow pixels due to the Cy3 and FITC fluorophores overlapping. However, in the *dpy-27 EQ* mutant background, the SDC-2 and DPY-27 puncta rarely, if ever colocalize (Figure 7A, row 2, far right).

**Figure 7.**
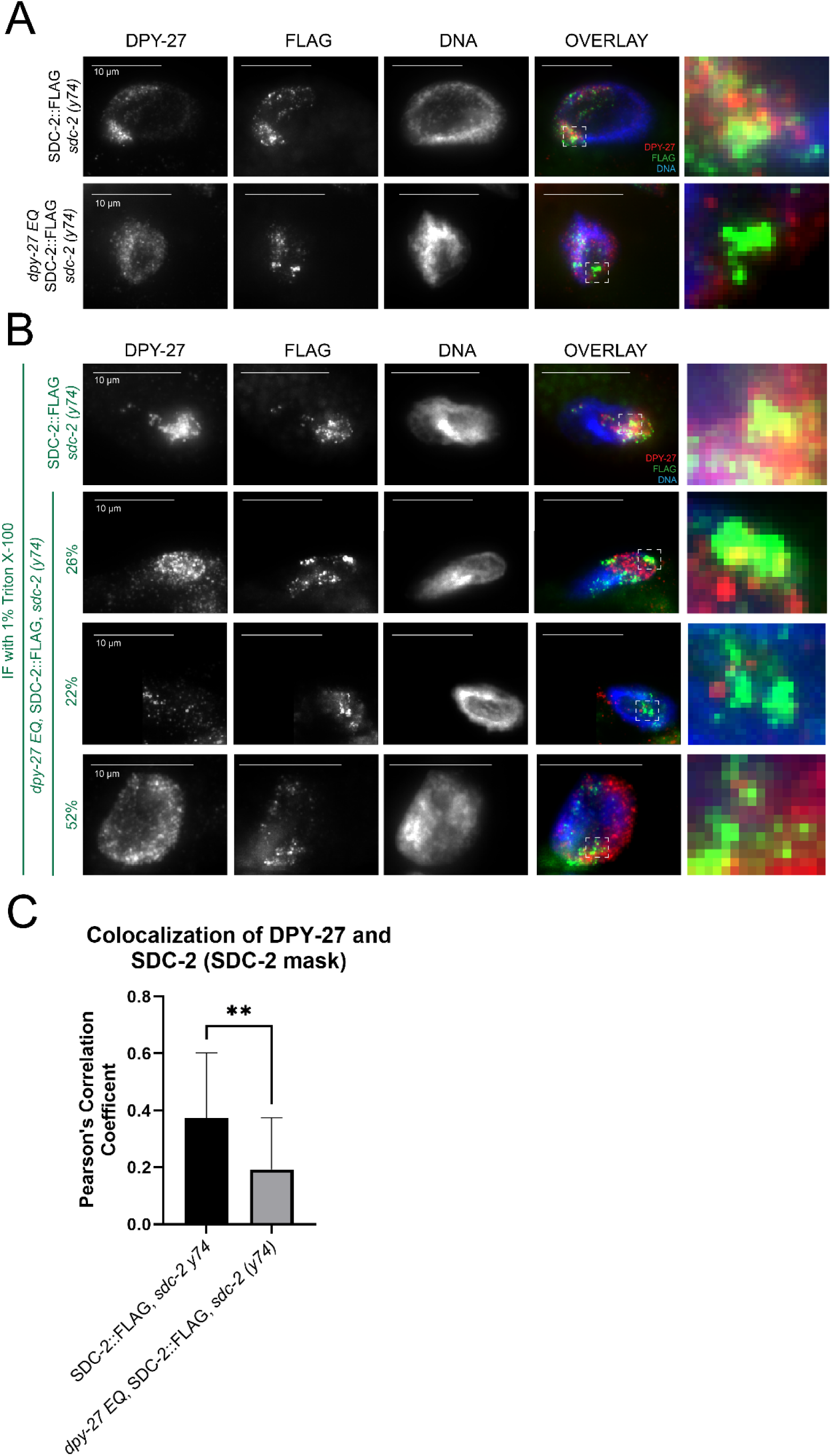
SDC-2 and DPY-27 localization in *dpy-27 EQ* mutants. **(A)** Using antibodies against FLAG and DPY-27, localization of SDC-2::FLAG and DPY-27 was assayed in gut nuclei of age-matched hermaphrodites. In SDC-2::FLAG, there is a high level of colocalization between SDC-2 and DPY-27. Colocalization is reduced in the *dpy-27 EQ* mutant (bottom row). Images in final column were created by zooming in and cropping images from the fourth column. All images are taken at the same magnification and exposure. **(B)** Same as (A) but with detergent extraction. In SDC-2::FLAG, DPY-27 and FLAG colocalize. When the *dpy-27 EQ* mutation is also present, some nuclei had restricted DPY-27 staining (26%, 7/27), some nuclei had no DPY-27 staining (22%, 6/27), and the rest of the nuclei had DPY-27 staining all over the nucleus (52%, 14/27). **(C)** Pearson’s correlation of DPY-27 and FLAG::SDC-2 within the FLAG::SDC-2 mask was measured in nuclei from (B). Quantification of *dpy-27 EQ* mutant nuclei was performed only in nuclei with real DPY-27 staining (2nd and final rows, B). Error bars indicate standard deviation. Statistical significance was determined by an unpaired student’s t-test with p < 0.0001.

To limit our analysis to strongly DNA-bound DPY-27 EQ protein only, we performed IF in these strains in the presence of detergent (Figure 7B). Similar to Figure 4B, we again saw three phenotypes of DPY-27 EQ protein: restricted staining (Figure 7B, 2nd row), no staining (Figure 7B, 3rd row), and staining all over the nucleus (Figure 7B, final row). We measured the Pearson’s correlation of DPY-27 and FLAG staining in nuclei with true DPY-27 signal and found that with the addition of the *dpy-27 EQ* mutation, there is significantly less correlation between DPY-27 and SDC-2 (Figure 7C). These results suggest that without ATPase function, condensin I^DC^ is unable to reach *rex* sites bound by SDC-2.

### ATPase function of DPY-27 is required for other DC processes

DC in *C. elegans* involves several mechanisms to accomplish the down regulation of each X-chromosome [73,84,96–98], including the enrichment of H4K20me1 on X chromosomes [67,97–100]. The *dpy-27 EQ* mutation produces a protein that is able to bind the X chromosome some of the time, and it colocalizes with another member of condensin I^DC^. Therefore, we asked if the DPY-27 EQ protein was still able to perform other DC associated functions. We decided to assay the enrichment of H4K20me1 to test this hypothesis. If DPY-27 is only a scaffold and ATPase function is not required for other processes, H4K20me1 should be enriched on X chromosomes in some *dpy-27 EQ* gut nuclei, at least in the subset of the nuclei in which the mutant protein was able to bind the X. Again, we combined the whole chromosome PAINT FISH and immunofluorescence to assay H4K20me1 enrichment (Figure 8A). In wildtype gut nuclei, H4K20me1 is clearly enriched on the X chromosomes. Interestingly, in all of the nuclei in both *dpy-27 EQ* and *dpy-27 Indel* worms, H4K20me1 signal is no longer enriched on the X chromosomes and instead dispersed throughout the nucleus, recapitulating other DCC mutants. These results suggest that ATPase function is required for H4K20me1 enrichment [98].

**Figure 8.**
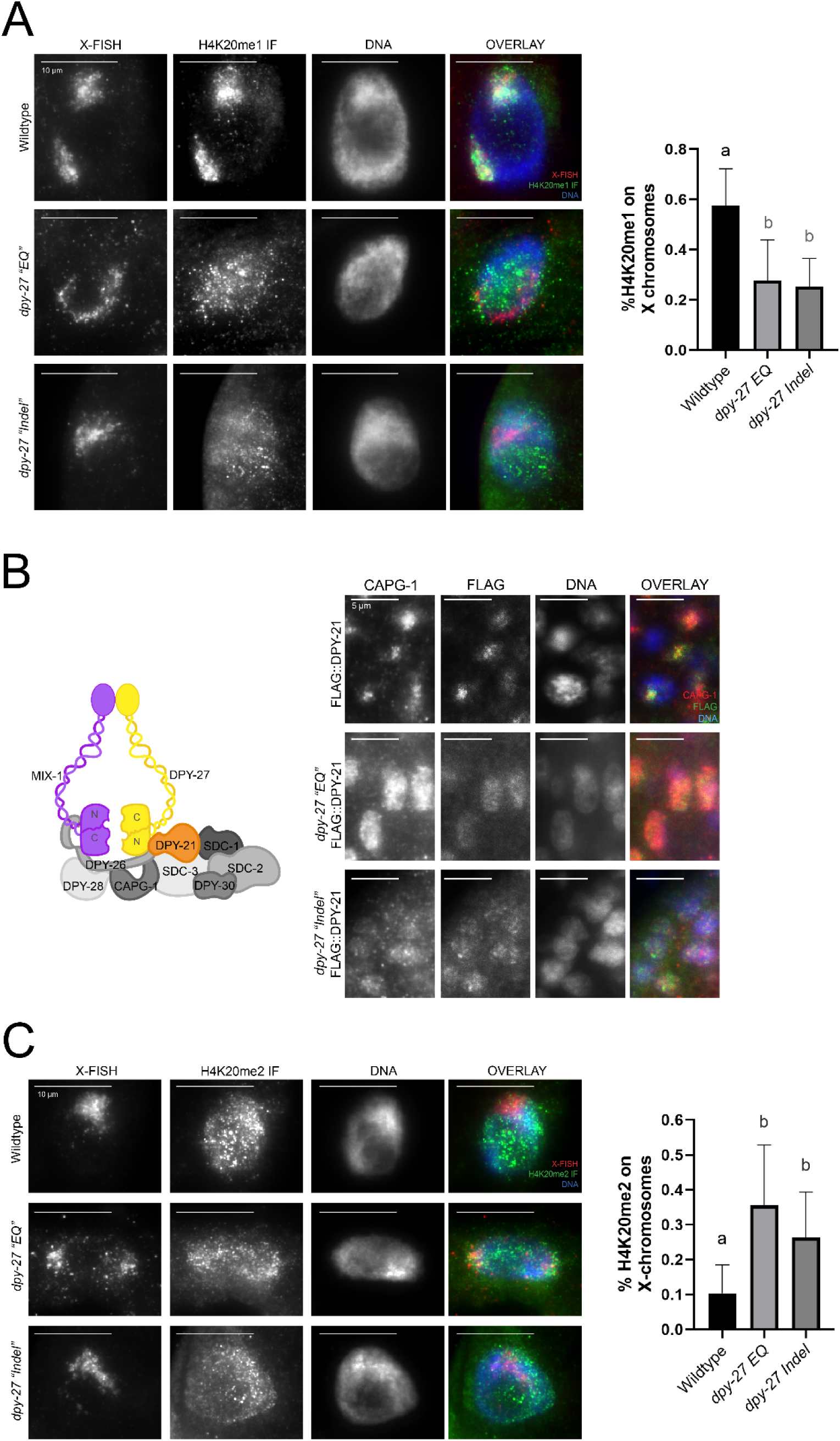
H4K20me state of *dpy-27* mutants. Using Whole X-PAINT FISH and antibodies, the methylation state of X chromosomes in age-matched differentiated gut nuclei was assayed. **(A)** In wildtype gut nuclei, H4K20me1 is specifically enriched on the X chromosomes (top), but this mark is no longer enriched only on the X chromosomes in either *dpy-27* mutant nuclei (second and bottom). Quantification shows percentage of H4K20me1 on X/total H4K20me1 signal. Error bars indicate standard deviation. Statistical significance was determined by a one-way ANOVA followed by Šídák’s multiple comparisons test. P-values are: N2 vs *dpy-EQ* p < 0.0001, N2 vs *dpy-27 Indel* p < 0.0001, *dpy-27 EQ* vs *dpy-27 Indel* p = 0.9229. **(B)** DPY-21 localization in dpy-27 mutant embryos. Using antibodies against FLAG and CAPG-1, DPY-21 localization was assayed in *dpy-27* mutants crossed into a FLAG::DPY-21 reporter line. In wildtype embryos, DPY-21 and CAPG-1 colocalize to the X chromosomes in nuclei at the bean stage (top). In *dpy-27 “EQ”* embryonic cells, DPY-21 and CAPG-1 colocalize but are diffusely spread over the entire nucleus in bean stage (middle). In *dpy-27 “Indel”* embryonic cells, DPY-21 and CAPG-1 staining is weak, suggesting that the DCC is not stable (bottom). **(C)** In wildtype gut nuclei, most of the nucleus has H4K20me2, but the signal is depleted on the X chromosomes (top). In both *dpy-27* mutant nuclei, the X chromosomes cannot be differentiated from the rest of the nucleus based on their H4K20me methylation state (middle and bottom). Quantification shows percentage of H4K20me2 on X/total H4K20me2 signal. Error bars indicate standard deviation. Statistical significance was determined by a one-way ANOVA followed by Šídák’s multiple comparisons test. P-values are: N2 vs *dpy-EQ* p < 0.0001, N2 vs *dpy-27 Indel* p = 0.0004, *dpy-27 EQ* vs *dpy-27 Indel* p = 0.0785.

H4K20me1 is produced by one of the non-condensin members of the DCC, DPY-21, a protein which associates with the X chromosomes after condensin I^DC^ [66,67]. To find out where DPY-21 is localized in our *dpy-27* mutants, we used FLAG::DPY-21, where the 3xFLAG tag has been added to the endogenous locus. DPY-21 is most highly expressed in embryos and there is little DPY-21 expression in adult worms [66], which we observed in preliminary experiments with the FLAG::DPY-21 version as well. Therefore, we performed these experiments in embryos. Using immunofluorescence, we stained for FLAG and CAPG-1, a condensin I^DC^ subunit (Figure 8B). In wildtype embryos, FLAG and CAPG-1 colocalize on the X chromosomes in each nucleus. In *dpy-27 EQ* embryos, we saw CAPG-1 staining over all of the nucleus, reminiscent of CAPG-1 staining in gut nuclei from Figure 4A. Interestingly, we also saw FLAG staining over the nucleus (Figure 8B, row 2).In contrast, there was very little CAPG-1 staining in *dpy-27 Indel* embryonic nuclei, again reminiscent of CAPG-1 staining in gut nuclei from Figure 4A. Interestingly, there was also very little FLAG staining, which suggests that FLAG::DPY-21 did not accumulate in the *dpy-27 Indel* embryonic nuclei the way it does in wildtype or *dpy-27 EQ* nuclei.

Enrichment of H4K20me1 occurs as a reduction from H4K20me2. The entire genome has H4K20me2 deposited by SET-4 [97], and the non-condensin DCC subunit DPY-21 demethylates this mark to H4K20me1 specifically on X chromosomes [67]. Therefore, we questioned whether the X chromosomes in *dpy-27* mutants had more H4K20me2, compared to wildtype nuclei where it has been depleted. Again, we combined the whole chromosome PAINT FISH and immunofluorescence to assay H4K20me2 depletion (Figure 8C). In wildtype gut nuclei, there is very little H4K20me2 signal on the X chromosomes. In both *dpy-27 EQ* and *dpy-27 Indel* mutants, there is H4K20me2 signal all over the nucleus, making it impossible to distinguish the X chromosomes from the autosomes based on methylation state. These results are consistent with DPY-21 not being able to localize to the X chromosomes and therefore not converting H4K20me2 to H4K20me1. Taken together, this data demonstrates that the ATPase function of DPY-27 is required to support other DC processes.

## Discussion

*C. elegans* contain an extra SMC protein, DPY-27, which functions within the DCC to equalize gene expression of the two hermaphrodite X chromosomes to the single male X. However, it was not known whether DPY-27 is capable of hydrolyzing ATP like other SMC proteins and if this function is required for DC. Using purified ATPase heads domains, we demonstrated that not only is DPY-27 capable of hydrolyzing ATP, but it accomplishes this at a rate comparable to its mitotic paralog, SMC-4. Moreover, we demonstrated that mutating the essential glutamate to a glutamine (EQ) in the Walker B domain abolishes ATPase function of DPY-27 *in vitro*. We then generated an ATPase dead mutant of *dpy-27 in vivo* and found that abolishing this function resulted in a loss of DC in homozygous mutant worms. The *dpy-27 EQ* mutation resulted in only a portion of the protein being able to bind DNA strongly, and only a subset localized to the X chromosomes. Even the portion of the mutant protein that appeared X localized, did not co-localize with SDC-2, the main recruiter of the DCC to the X. The EQ mutation also resulted in decondensed X chromosomes and lack of H4K20me1 enrichment, indicating that many DC related processes are disrupted. Due to their role in mitosis, many studies on condensin’s gene regulatory role are limited to using cell viability as a functional readout [47,101], limited to a subset of post-mitotic cells [52,53], using *ex vivo* systems [13,36], or are entirely *in vitro* [26,37,102]. By generating an ATPase dead mutant of DPY-27 and demonstrating that this function is required for DC, we have created an *in vivo* model to study how condensin’s enzymatic function impacts gene regulation.

### DPY-27 is an SMC protein capable of hydrolyzing ATP

Previous work has frequently used DPY-27 to study the function of condensin I^DC^ as it is the only unique member of the complex [16], but only a few studies attempted to understand how DPY-27 functions as an SMC protein [77]. DPY-27 shares a high degree of homology with other SMC proteins, and therefore, mechanistic studies of those other SMC proteins [13,15,18,103], have been suggested to apply to DPY-27. For example, findings in other organisms showed that condensins are capable of extruding DNA loops, an activity that can lead to the formation of TADs [104,105]. In *C. elegans*, the X chromosomes have clearly visible TADs, similar to TADs in other organisms, and their formation is the result of DCC and condensin I^DC^ function [73]. Autosomal TADs are generally much smaller and mostly dependent on histone modifications to define their boundaries [73,74,100,106]. The dependence of TAD formation on condensin I^DC^ mirrors how cohesin function is required for TAD formation in other species [104,107–110]. Here, our work adds context by demonstrating that DPY-27 actively uses ATP, functioning like a true SMC protein. This finding also allows us to now consider that DPY-27 and condensin I^DC^ may be capable of loop-extrusion, as it has been demonstrated *in vitro* by other condensins and cohesins [75,102,107,109,111], and that this loop extrusion function is the true source of TADs, as seen in other organisms [109]. We also demonstrated that without ATPase function, *dpy-27 EQ* mutants had decondensed X chromosomes (Figure 6). While the X chromosomes can still condense when TAD formation is reduced after deletions of some TAD boundaries [112], we know that both condensation and TAD formation require functional DPY-27 and therefore we can predict that *dpy-27 EQ* mutants will have weaker TADs compared to wildtype.

We also compared *dpy-27 EQ* heterozygous and homozygous worms to check if the mutation behaved as a dominant negative. Some proteins that oligomerize have been shown to become dominant negative when mutated, where the mutant protein will poison the wildtype [113]. Dominant negative mutants can also interfere with complex formation [114,115], or entirely complex function [116]. As DPY-27 works in condensin I^DC^ and must dimerize with MIX-1, we questioned whether worms containing both wildtype and EQ mutant DPY-27 protein would result in a similar phenotype because condensin I^DC^ containing DPY-27 EQ protein would prevent functioning of wildtype condensin I^DC^. Interestingly, we observed that in *dpy-27 EQ/+* heterozygous worms, the mutant form of the protein did not seem to impede the function of wildtype DPY-27 (Figure 2, Supplemental Figure 4). One possible explanation is that condensin I^DC^ with wildtype DPY-27 may be able to transverse past DPY-27 EQ containing condensin I^DC^, as other condensins have been demonstrated capable of *in vitro* [117].

### The ATPase function of DPY-27 affects DNA binding

Some *in vitro* studies have shown that loss of ATPase ability does not prevent *S. cerevisiae* condensin from binding DNA but prevents it from continuously loop extruding DNA [111]. By contrast, studies from *S. pombe* and *B. subtilis* demonstrated that loss of ATPase function has an impact on both functions [51,118]. Single-molecule experiments have attempted to describe the mechanism of loop extrusion and ATP hydrolysis, and one mechanism states that the SMC heads are kept apart by the kleisin. However, when ATP is bound, the kleisin-SMC head interaction weakens and the kleisin moves, allowing for DNA to enter the “ring” between the SMC protein α-helices. DNA is trapped once the SMC heads hydrolyze ATP and return to binding the kleisin [51,119,120]. By combining the *in vitro* condensin data with our observations, we can hypothesize that the stably DNA-bound DPY-27 foci are DPY-27 protein that bound DNA after binding ATP because it opened the SMC-kleisin ring. With the ATPase dead mutation, these molecules may be trapped in the “transition state” [51] as they cannot hydrolyze the ATP molecules, and therefore strongly bound to DNA (Figure 9A). In contrast, weakly DNA-bound DPY-27 foci are DPY-27 protein that entered the transition state before DNA could enter the SMC-kleisin ring and therefore are only DNA-associated through DNA interactions by the non-SMC subunits of condensin I^DC^.

**Figure 9.**
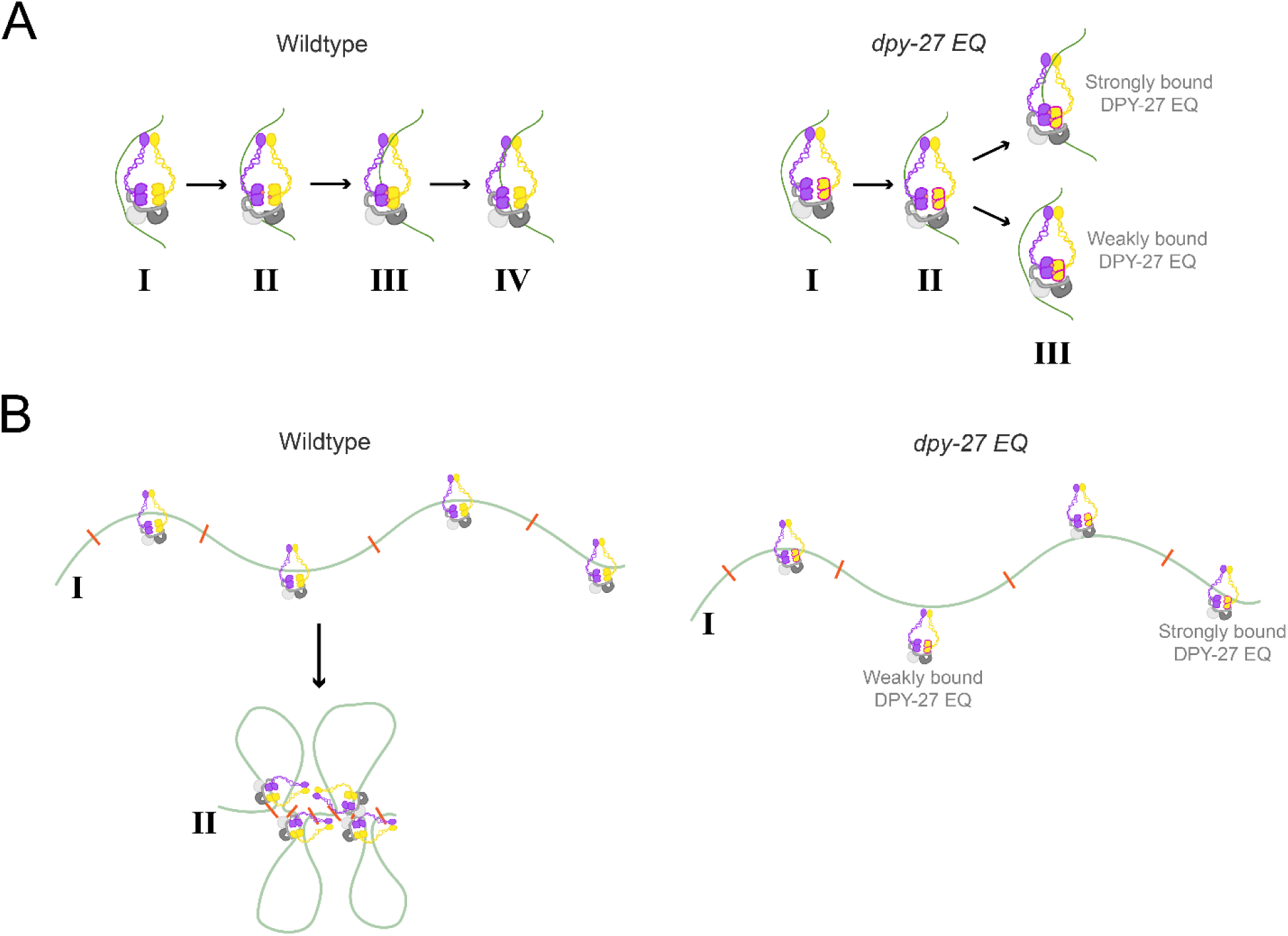
Model of ATP-dependent loop extrusion and DNA binding. **(A)** Wildtype condensins are thought to bind DNA through an ATP-dependent mechanism as follows: (I) condensin I^DC^ associates with DNA through the SMC hinges and HAWKs, (II) the SMC heads bind ATP, which opens the ring and allows DNA to enter, (III) the SMC heads come together to hydrolyze ATP, and (IV) after hydrolyzing ATP, the SMC heads return to their original conformation, which closes the ring and traps the DNA within it, making condensin I^DC^ “strongly DNA-bound”. When DPY-27 EQ is present, the first steps should remain the same, with condensin I^DC^ associating with DNA through the SMC hinges and HAWKs (I), and the SMC heads opening the ring after binding DNA. However, when the heads come together to hydrolyze ATP (III), they get trapped in a “transition state” [51] because of the *EQ* mutation. If DNA is inside the ring before the heads are trapped, then condensin I^DC^ is strongly DNA-bound. If DNA is not inside the ring before the heads get trapped, then condensin I^DC^ is weakly DNA-bound. **(B)** In wildtype nuclei, condensin I^DC^ binds to DNA randomly **(I)**. Then, it loop-extrudes until it halted at *rex* sites (orange dashes) **(II)**. In *dpy-27 EQ* nuclei, condensin I^DC^ randomly binds DNA either strongly or weakly **(I)**, but cannot loop-extrude chromatin to get to rex sites without ATPase function.

An interesting question this work brings up is how the DCC binds and stays on the X chromosomes. Previous work in the field has demonstrated that DCC binds most strongly at *recruitment element on X (rex)* sites along the X chromosomes and less strongly at promoters of genes [73,92–95,121]. However, the mechanism has been debated. One pool of evidence supports a “spreading model” where the entire DCC is initially recruited to *rex* sites before spreading to additional sites to cover the entirety of X chromosomes [74,93,122,123]. The alternative “roadblock model” suggests the idea that condensin I^DC^ initially binds to random sites on the chromosomes then translocates along DNA until it is stopped at *rex* sites by SDC-2 [106,112]. Part of our data supports the second model. In Figures 4, 6, and 9, we observed that DPY-27 EQ protein binds DNA randomly throughout the nucleus and does not colocalize with SDC-2. From previous literature, we know that condensins in other organisms have no sequence specificity when it comes to binding DNA and rely on other proteins to anchor them to specific regions [124], similar to how CTCF anchors cohesin [125]. Moreover, condensins in other organisms require ATP hydrolysis to loop extrude and translocate DNA [102,111]. It is possible that DPY-27, like other condensins, loads onto DNA randomly and the high degree of colocalization between SDC-2 and wild type DPY-27 results from DPY-27 loop extruding DNA until it reaches SDC-2-bound rex sites. DPY-27 EQ protein cannot hydrolyze ATP. Therefore, we can hypothesize that the decreased DPY-27 EQ localization with SDC-2 implies that the DPY-27 EQ cannot loop-extrude along DNA to *rex* sites from where it initially bound (see model in Figure 9B). Ideally, a detailed study of DPY-27 EQ binding at high resolution would provide more evidence to support this. Unfortunately, due to the maternal effect lethal phenotype in *dpy-27 EQ* worms and the need to maintain the strain as a heterozygote, isolating a pure population of mutant worms for these localization experiments is not possible to achieve with the reagents on hand.

### Unexpected phenotypes in dpy-27 EQ mutants

While it was not statistically significant compared to wildtype, there was increased embryonic lethality in the progeny of both *dpy-27 EQ/+* and *dpy-27 Indel/+* heterozygous worms (Figure 3). One explanation could be that with heterozygous worms, there is random loading of both wildtype and mutant protein into the embryos. Taking the zygotic rescue assay into account, the increased embryonic lethality among the progeny of the heterozygotes could be of homozygous mutant embryos that received more mutant protein than wildtype and also could not produce wildtype DPY-27 protein themselves to prevent larval arrest.

Previous literature has demonstrated that DPY-21 is dependent on DPY-27 to reach the X chromosomes [66], and we saw that on average, 35% of DPY-27 EQ protein did reach the X chromosomes (Figure 6, Supplemental Figure 3). Therefore, we wondered if there would be partial enrichment of H4K20me1 on the X chromosomes where DPY-27 EQ was bound. To our surprise, we observed that there was no discernable enrichment of H4K20me1, even though there is discernable enrichment of DPY-27 EQ on the X chromosomes in the *dpy-27 EQ* mutants (Figure 8A). Furthermore, DPY-21 was found all over the nucleus in *dpy-27 EQ* mutants (Figure 8B). Therefore, our results demonstrate that recruitment of DPY-21 to the DCC, and subsequently to the X chromosomes, is dependent on DPY-27 being able to hydrolyze ATP.

In contrast, *dpy-27 Indel* mutants having no DPY-21 staining in the nucleus is unexpected. Previous literature has demonstrated that DPY-21 only needs other DCC members for localization to the X chromosomes, not for protein stability [66]. We assayed DPY-21 localization in embryos at a later developmental stage than the previous studies. DPY-21 has been shown to be expressed at high levels in embryogenesis, but this expression decreases quickly as the embryo develops and becomes minimal in hatched worms [66]. Previously published literature has observed DPY-21 localization in pachytene nuclei in the germline [126] or 40-cell or younger embryos [66]. While DPY-21 can localize as early as the 40-cell stage [66], enrichment of H4K20me1 happens after the 100-cell stage [127]. We used the embryos closer to the 200-cell stage as we wanted to use a developmental stage that was well after DPY-21 should be localized and functioning on the X chromosomes. Therefore, it is possible that at this later developmental stage, DPY-21 does not remain in the nucleus without DPY-27.

### C. elegans DC as a model for studying condensin function in chromosome-wide gene regulation

The requirement of ATP for condensin-mediated compaction has been shown in multiple *in vitro* studies [39,101,111]. As we observed in Figure 6, *dpy-27 EQ* mutants had severely decondensed X chromosomes. This differs from mitotic condensin defects *in vivo*, where chromosomes are still able to condense but instead have defects in segregation [43,128]. One possible explanation could be that mitotic condensation has multiple drivers of equal consequence including changes in histone modifications [129], whereas condensation of dosage compensated X chromosomes is predominantly dependent on the DCC.

On a broader scale, the *C. elegans* DCC allows for the dissection of how condensins work with other mechanisms. For example, work in *C. elegans* has demonstrated how topoisomerases work with condensin I^DC^ to regulate X-chromosome gene regulation [130], building on work on the interaction between condensins and topoisomerases from dividing yeast cells [131]. Using *C. elegans* condensin I^DC^ as a model for other condensins also allows for the dissection of condensin-mediated gene regulation over the course of development. Previous studies found that condensins are responsible for heterochromatin formation in *D. melanogaster* neurons, but this was done in post-mitotic neurons [52]. Now with the DCC, we can ask how condensin-mediated gene expression occurs throughout development and apply these findings more broadly.

Taken together, our work helps bridge the gap between the *in vitro*, *ex vivo*, and *in vivo* studies on SMC protein function. By demonstrating that DPY-27 is a true SMC protein and condensin I^DC^ acts similar to other condensins, this allows us to now dissect the ways in which condensins mediate gene regulation separately from their role in cell division.

## Author Contributions

Conceptualization, B.C. and G.C.; Experimentation: cloning, protein purification, and biochemistry: B.C. and S.J.; CRISPR design and injection: D.S. and G.C.; *C. elegans* experimentation and analysis: B.C.; Writing—original draft preparation, B.C.; Writing— review and editing, B.C. and G.C.; Supervision, G.C.; Project Administration, G.C.; Funding Acquisition, G.C. All authors have read and agreed to the published version of the manuscript.

## Acknowledgments

This research was supported by the National Institutes of Health, grant numbers R01GM13385801 (to G.C.) and R35GM149543 (to G.C.). Some strains were obtained from the Caenorhabditis Genetics Center which is funded by NIH Office of Research Infrastructure Programs (P40 OD010440). We thank JK Nandakumar for sharing cloning plasmids as well as access and usage of his FPLC, as well as members of his lab for training and supervision.

