## Supplemental Figures for "Condensin I^DC^ has a Functional ATPase That is Required for X-Chromosome Dosage Compensation in *C. elegans*"

| | N-term | A-loop | P-loop | R-loop | Q-loop |
|---|---|---|---|---|---|
| SMC-1 isoform a | | ENFKSYK | IGPNGSGKSNL | SLRV | VYQGAI |
| SMC-3 isoform a | | TGFRSYK | VGRNGSGKSNF | HLKE | VKQGKI |
| MIX-1 | | DGFKSYQ | TGYNGSGKSNI | NIRA | IMQGRI |
| SMC-4 | | DNFKSYF | IGPNGSGKSNL | KIRS | ILQGEV |
| DPY-27 | | ENFKSYA | LGPNGSGKSNV | KIRT | ILQGEV |
| *C. thermophilum* Smc4 | | TNFKSYA | VGPNGSGKSNV | KMRQ | ILQGEV |
| *S. pombe* Cut3 | | TNFKSYA | VGPNGSGKSNV | KLRQ | ILQGEV |
| *S. cerevisiae* SMC4 | | ENFKSYA | VGPNGSGKSNV | KMRQ | ILQGEV |
| *Human* SMC4 | | QNFKSYA | IGPNGSGKSNV | KIRS | ILQGEV |
| *X. laevis* SMC-4 | | QNFKSYA | IGPNGSGKSNV | KIRS | ILQGEV |
| *P. furiosus* Smc | | KGFKSYG | VGANGSGKSNI | AMRA | VLQGDI |
| *B. subtilis* Smc | | IGFKSFA | VGPNGSGKSNI | SLRG | ISQGKV |
| *X. laevis* SMC-2 | | DGFKSYA | TGLNGSGKSNI | QVRA | IMQGRI |
| *P. furiosus* Rad50 | | KNFRSHS | IGQNGSGKSSL | RIKD | IRQGQI |
| | | .*.*. | * ******.. | :. | . ** . |
| *Human* MDR1 | | VHFSYPS | VGNNGSGKSTT | DIRT | VSQEPV |
| *Human* CFTR | | LFFSNFS | AGSTGAGKTSL | KIKH | CSQFSW |

| | C-motif | C-helix | Walker B | D-loop | H-loop | C-term |
|---|---|---|---|---|---|---|
| SMC-1 isoform a | LSGGEK | ALALL | VLDEI | DAALD | QIIVISL | |
| SMC-3 isoform a | LSGGQK | ALAII | LFDEI | DAALD | QFVTTTF | |
| MIX-1 | LSGGQR | ALSLI | ILDEV | DAALD | QFIIVSL | |
| SMC-4 | LSGGEK | SLALI | VMDEI | DAALD | QFIIISL | |
| DPY-27 | LSGGEK | SLCFV | VMDEI | DAALD | QFIIISL | |
| *C. thermophilum* Smc4 | LSGGEK | SLALV | VMDEI | DAALD | QFIVISL | |
| *S. pombe* Cut3 | LSGGEK | SLALV | VMDEI | DAALD | QFIVISL | |
| *S. cerevisiae* SMC4 | LSGGEK | SLALV | VMDEI | DAALD | QFIVISL | |
| *Human* SMC4 | LSGGEK | SLALV | FMDEI | DAALD | QFIIISL | |
| *X. laevis* SMC-4 | LSGGEK | SLALV | FMDEI | DAALD | QFIIISL | |
| *P. furiosus* Smc | MSGGEK | ALAFV | LFDEI | DAHLD | QFIVITL | |
| *B. subtilis* Smc | LSGGER | AIALL | VLDEV | EAALD | QFIVITH | |
| *X. laevis* SMC-2 | LSGGQR | ALSLI | ILDEV | DAALD | QFIVVSL | |
| *P. furiosus* Rad50 | LSGGER | GLAFR | ILDEP | TPYLD | QVILVSH | |
|  | :***:. | .:.: | .:** | . ** | *.: : |  |
| *Human* MDR1 | LSGGQK | AIARA | LLDEA | TSALD | TCIVIAH | |
| *Human* CFTR | LSHGHK | CLARS | LLDEP | SAHLD | TVILCEH | |

**Supplemental Figure 1 Sequence Alignments of SMC protein ATPase head domains.**The top row consists of the ATPase residues found in the N-terminal globular domain while the bottom row consists of those residues found in the C-terminal globular domain. Sequences of SMC proteins found in *C. elegans* are in shades of green. SMC-4 paralogs from species other than *C. elegans* are in blue. Sequences of SMC proteins other than SMC-4 from species other than C. elegans are purple. ABC transporter protein sequences are orange. Residues with perfect identity throughout all proteins are in red and have a star at the bottom of the column. Residues with near perfect identity throughout all proteins are in black and have a dot at the bottom of the column. Residues with highly similar identity throughout all proteins are black and italicized, with a colon at the bottom of the column. Scores do not include the ABC transporters.

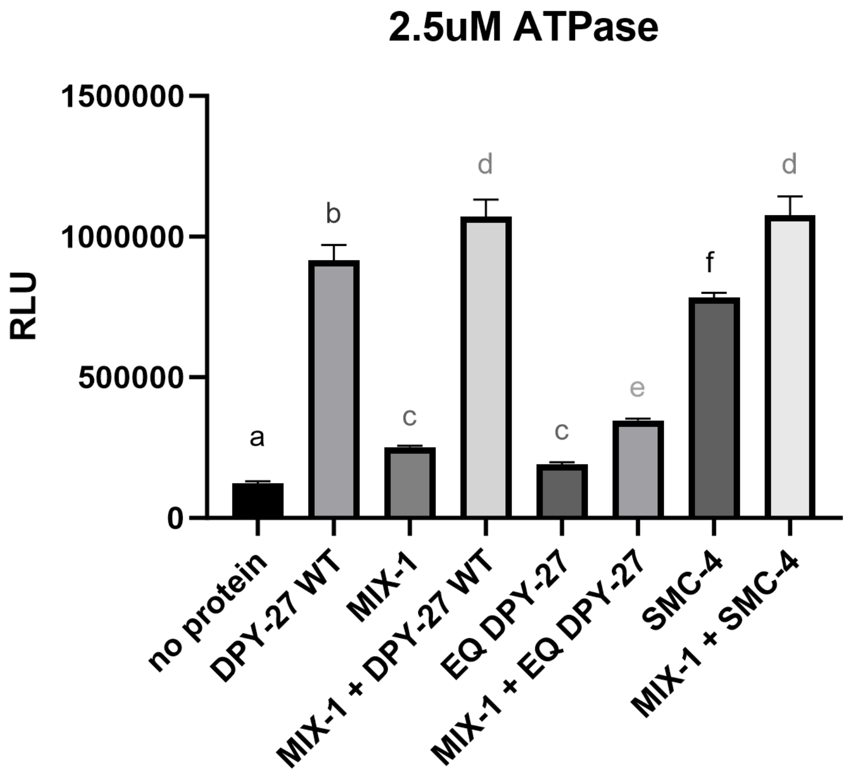


**Supplemental Figure 2 *In Vitro* ATPase Assay with single proteins.**Using the Promega ADP-Glo Kit, the rate of ATP hydrolysis was assayed for the different SMC "ATPase head" proteins, either as heterodimers or as individual proteins. The heterodimer combinations of MIX-1 + DPY-27 WT and MIX-1 + SMC-4 produced the most ADP in the 15-minute reaction time. The monomers of DPY-27 WT or SMC-4 were also capable of hydrolyzing ATP at a high level, possibly by forming homodimers. In contrast, the homodimers of MIX-1 or EQ DPY-27 were barely able to hydrolyze ATP over the basal rate of ATP degradation in solution (no protein). Interestingly, the heterodimeric combination of MIX-1 + EQ DPY-27 hydrolyzed more ATP than the homodimeric reactions, demonstrating that the heterodimeric combination is more efficient and that the MIX-1 walker B is functional. 3 replicates of each reaction were run and a one-way ANOVA with multiple comparisons was used to determine statistical significance. Letters represent multiple comparison p values, with different letters indicating statistically significant differences, and any repeated letter demonstrating no statistically significant difference. See Supplemental Table 4 for all p-values.

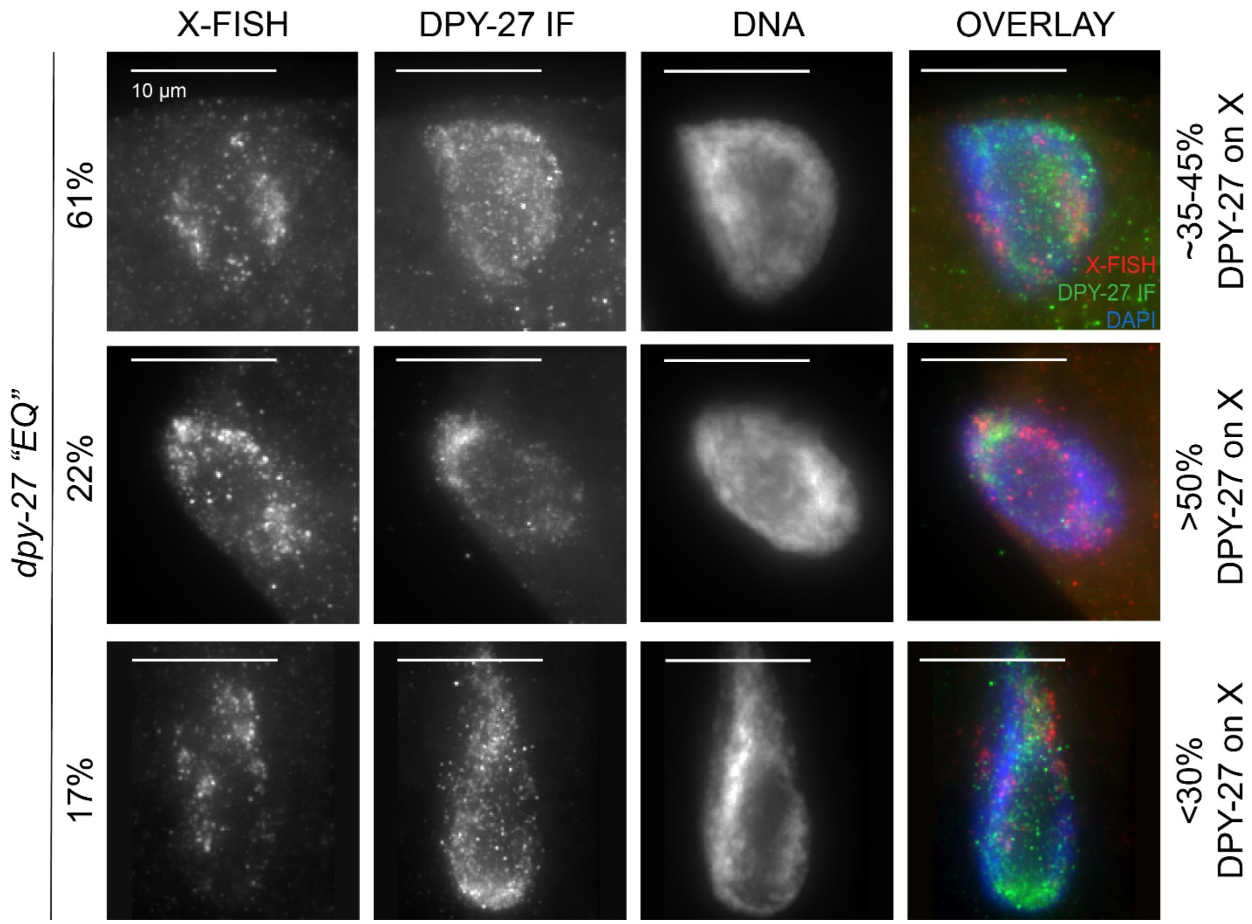


**Supplemental Figure 3 DPY-27 EQ protein phenotypes and X-localization.** DPY-27 EQ phenotypes were binned into phenotypes based on percentage of DPY-27 signal on the X chromosomes. Most nuclei were in the first category, where DPY-27 signal is dispersed through the nucleus, corresponding to only 35-45% of the DPY-27 signal being on the X chromosomes. The second phenotype was characterized by 50% or more DPY-27 signal on the X chromosomes, but the entire X chromosomes were not covered by DPY-27 signal. The final phenotype was characterized by dispersed DPY-27 signal all over the nucleus, but less than 30% of the signal on the X chromosomes. n numbers for each category are: 14/23, 5/23, and 4/23, respectively.

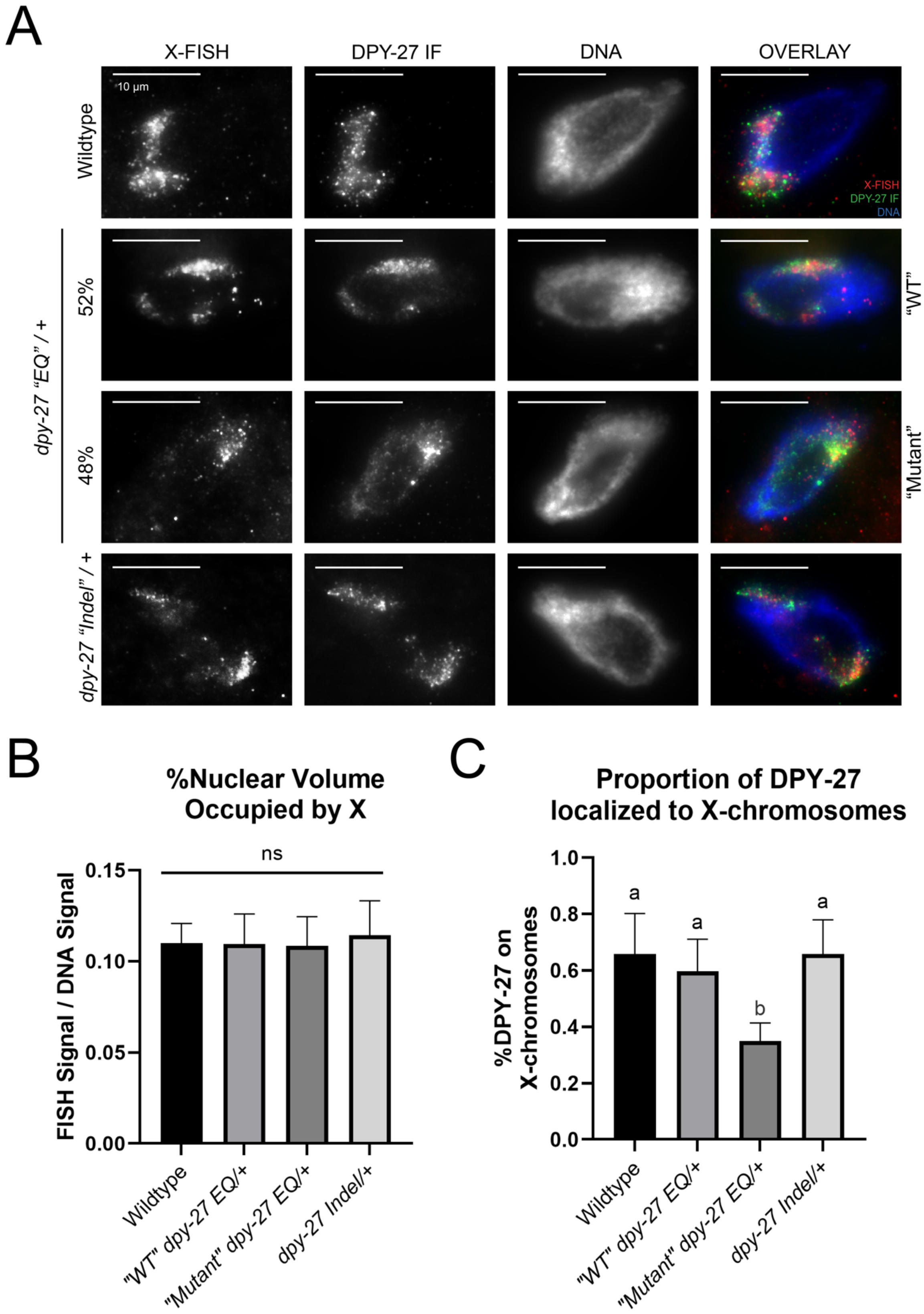
A
X-FISH
DPY-27 IF
DNA
OVERLAY
10 µm
Wildtype
X-FISH
DPY-27 IF
DNA
dpy-27 "EQ" / +
52%
48%
"WT"
"Mutant"
dpy-27 "Indel" / +
B
%Nuclear Volume Occupied by X
FISH Signal / DNA Signal
ns
0.15
0.10
0.05
0.00
Wildtype
"WT" dpy-27 EQ/+
"Mutant" dpy-27 EQ/+
dpy-27 Indel/+
C
Proportion of DPY-27 localized to X-chromosomes
%DPY-27 on X-chromosomes
1.0
0.8
0.6
0.4
0.2
0.0
a
a
b
a
Wildtype
"WT" dpy-27 EQ/+
"Mutant" dpy-27 EQ/+
dpy-27 Indel/+

**Supplemental Figure 4 DPY-27 localization on X chromosomes in heterozygous *dpy-27* worm strains. (A)** Using Whole X-PAINT FISH and antibodies against DPY-27, localization of DPY-27 in differentiated gut nuclei of age-matched hermaphrodite worms was assayed. In wildtype nuclei, DPY-27 is found on the X chromosomes (row 1). *dpy-27 Indel/+* heterozygous nuclei phenocopy wildtype nuclei, with DPY-27 found on the X-chromosomes (row 4). In *dpy-27 EQ/+* heterozygotes, two phenotypes of nuclei were found. In half of the nuclei, most of the DPY-27 signal was on the X chromosomes, resembling wildtype (denoted “WT”). In the other half, there was a proportion of DPY-27 signal on the X chromosomes, but also significant staining elsewhere in the nucleus, resembling dpy-27 EQ homozygous mutant nuclei (denoted “Mutant”). **(B)** Quantification of X-chromosome nuclear occupancy. Error bars indicate standard deviation. Statistical significance was determined by a one-way ANOVA ($p = 0.639$). **(C)** Quantification of DPY-27 on X-chromosomes. Error bars indicate standard deviation. Statistical significance was determined by a Brown-Forsythe ANOVA ($p < 0.0001$) and letters represent Dunnett’s T3 multiple comparison p values, with any repeated letter demonstrating no significance. See Supplemental Table 6 for p values.

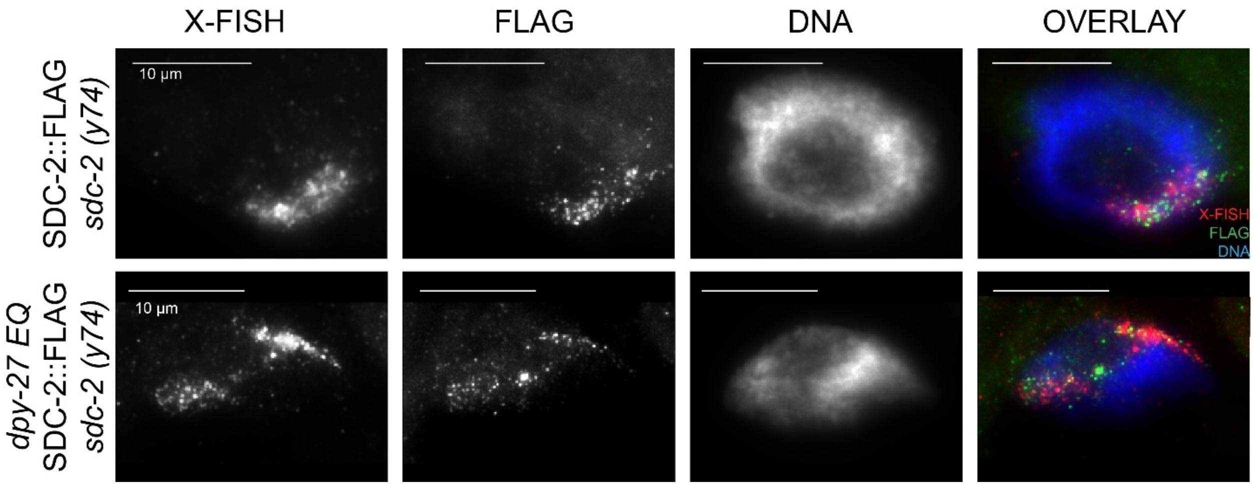


**Supplemental Figure 5 SDC-2 localization in *dpy-27 EQ* mutants.** Using probes for the X chromosomes and antibodies against FLAG in SDC-2::FLAG, *sdc-2 (y74)* background (top row), FLAG staining demonstrates that SDC-2 localizes to the X chromosomes as expected. In the *dpy-27 EQ* background (bottom row), FLAG staining demonstrates that SDC-2 still localizes to the X-chromosomes even when DPY-27 is mutant.
