## Supplemental Tables for "Condensin I^DC^ has a Functional ATPase That is Required for X-Chromosome Dosage Compensation in *C. elegans*"

**Supplemental Table 1 Protein Sequences used for alignments.**

| Protein | *Organism* | Identifier | Source |
|---|---|---|---|
| HIM-1 (SMC-1) isoform a | *C. elegans* | F28B3.7a | Wormbase |
| SMC-3 isoform a | *C. elegans* | Y47D3A.26a | Wormbase |
| MIX-1 | *C. elegans* | M106.1 | Wormbase |
| SMC-4 | *C. elegans* | F35G12.8 | Wormbase |
| DPY-27 | *C. elegans* | R13G10.1 | Wormbase |
| Smc4 | *C. thermophilium* | G0S2G2 | Uniprot |
| Cut3 | *S. pombe* | P41004.2 | Uniprot |
| SMC4 | *S. cerevisiae* | NP_013187.1 | Uniprot |
| SMC4 | Human | NP_001002800.1 | Uniprot |
| SMC4 | *X. laevis* | NP_001081371.1 | Uniprot |
| Smc | *P. furiosus* | WP_014835559.1 | Uniprot |
| Smc | *B. subtilis* | NP_389476.2 | Uniprot |
| SMC2 | *X. laevis* | NP_001081372.1 | Uniprot |
| Rad50 | *P. furiosus* | P58301.1 | Uniprot |
| MDR1 | Human | NP_001335875.1 | Uniprot |
| CFTR1 | Human | P13569.3 | Uniprot |

### Supplemental Table 2 DNA Sequences Used in this study

| DNA Sequence Name | Sequence |
|---|---|
| **DPY-27 N-term ATPase Head (codon-optimized for *E. coli*)** | ATGATAATTCTGAATATCTACGTTGAGAACTTTAAAAGTTATGCTGGAAAA<br>CATATTCTCGGACCGTTTCACAAGAACTTGACAATGATCCTGGGACCAAAT<br>GGAAGTGGAAAGTCAAACGTTATTGATGCATTGCTATTCGTGTTTGGATTT<br>AAAGCTGGAAAAATTCGTACCAAGAAGCTATCAGCATTGATCAATTCAGG<br>AGGGAACTACGAATCATGTTCAGTAACAATTATGTTCCAGATGGTGAAGGA<br>TATGCCTGTCGAAAATTACGATAAATATGAAGTGTTAACAGATAATTGCGTC<br>TGCATCACCCGCACAATAAATCGTGAAAACAATTCAAAATATCGTATTGAT<br>GACAAAGATGCATCGCAAAAAGACGTCCAGGAGCTTCTCCTACGTGCTGG<br>AATTGATATGACCCATAATCGATTCCTGATTCTGCAAGGAGAAGTTGAAGC<br>GATTGCACTAATGAAGCCAACATCCAAAAATCCGAATGAAGAAGGAATGC<br>TTGAATATATTGAGGATATTGTTGGAACCAATCGTTTTGTCGCACCAATTTC<br>CAAACTAATGCAT |
| **DPY-27 C-term WT ATPase Head (codon-optimized for *E. coli*)** | ATGGCGAGATTCAACGAGTTCAGCGAGGCGCTCGCATTCTTGGGTACTAC<br>TACCCAAATGCTTTACCAGCTTATCACCAATGGCGGAGATGCTAGTTTGAA<br>GTTTGTGGAAGAAGGAAAATCTACCGATCCCTTCGATGGTGGAATAAAGT<br>TCAGTGTACGTCCCGCCAAGAAATCCTGGAAGCTCATCGAGAATCTATCTG<br>GCGGAGAGAAGACTCTGGCCTCTTTATGCTTTGTCTTTGCAATGCACCACT<br>ACCGTCCAACACCCCTCTACGTGATGGATGAAATCGATGCGGCACTGGAC<br>TTGAACAATGTCAGCCTGATTGCAAACTATATCAAGCATTCCGAGCGAACA<br>CGGAACGCTCAATTTATCATAATTTCGCTTCGAAATCAAATGTTCGAGGTC<br>GGAAATCGCTTGCTTGGTATCTATAAAATCGATGGAAAAACTTATAACATTA<br>TGGTGGATCCGATCGCGGTGGAGATCAAGAATCGTCCGATTTTGAAGATTT<br>TCGAGGAGGAGATCAAACGGCGCGAGAAG |
| **DPY-27 C-term EQ ATPase Head (codon-optimized for *E. coli*)** | ATGGCGAGATTCAACGAGTTCAGCGAGGCGCTCGCATTCTTGGGTACTAC<br>TACCCAAATGCTTTACCAGCTTATCACCAATGGCGGAGATGCTAGTTTGAA<br>GTTTGTGGAAGAAGGAAAATCTACCGATCCCTTCGATGGTGGAATAAAGT<br>TCAGTGTACGTCCCGCCAAGAAATCCTGGAAGCTCATCGAGAATCTATCTG<br>GCGGAGAGAAGACTCTGGCCTCTTTATGCTTTGTCTTTGCAATGCACCACT<br>ACCGTCCAACACCCCTCTACGTGATGGATCAAATCGATGCGGCACTGGAC<br>TTGAACAATGTCAGCCTGATTGCAAACTATATCAAGCATTCCGAGCGAACA<br>CGGAACGCTCAATTTATCATAATTTCGCTTCGAAATCAAATGTTCGAGGTC<br>GGAAATCGCTTGCTTGGTATCTATAAAATCGATGGAAAAACTTATAACATTA<br>TGGTGGATCCGATCGCGGTGGAGATCAAGAATCGTCCGATTTTGAAGATTT<br>TCGAGGAGGAGATCAAACGGCGCGAGAAG |
| **MIX-1 N-term ATPase Head (codon-optimized for *E. coli*)** | ATGCACATTAAATCCATCCATCTGGATGGCTTTAAATCGTACCAAAAGCAC<br>ACCGATATTCTCGATTTTTCGCCGACTTTCAACGCAATCACCGGATACAAT<br>GGTAGTGGAAAATCCAATATTCTCGATTCGATCTGCTTCATTATGGGAATCA<br>ATAAGCTCGATAATATCAGAGCAAAATCAATGCACGAGCTGATTTCCCATG<br>GTGGAACAAAAGCGATCGTCCAAGTTCGCTTCGACAACACTGATAAAAGA<br>TGCTCCCCGTTCGGAATGGAACATCTCGACGAGATCGTCGTTCAGAGAAT<br>CATCACAGCTCAAGCAACTGGAAAAGGATGTGCAACGAGTTACACACTG<br>AATGGGCATGCGGCCACAAATGGAAAAATGCAAGATTTCTTCCGTGGCGT<br>CGGTTTGAACGTGAATAATCCACATTTTCTCATCATGCAAGGTCGTATTACT<br>ACGGTGCTAAACATGAAGCCGGAGGAAATCCTCGGAATGGTTGAGGAAG<br>CAGCTGGAACAAAGATGTATGATCAGAAG |
| **MIX-1 C-term ATPase Head (codon-optimized for *E. coli*)** | AAAGTGGACGAGTTGATTAGGGCACACGAGTCGGTGAACAAGGACTTTG<br>GACAAATCTTCAACTGCCTGCTTCCCGATGCCCATGCCTCACTGGTACCTC<br>CAGAGGGAAAGACAGTTTGTGAGGGTCTTGAAGTGAAAGTCTCATTCGGT<br>GGAGTCGTCAAGGATAGTTTACATGAACTTTCTGGAGGACAGCGGTCTCT<br>TGTCGCGTTGTCACTGATTTTAGCTATGCTGAAATTCAAGCCAGCTCCATT<br>GTATATTCTCGACGAAGTTGATGCGGCCCTTGATTTGTCACACACCGCGAA<br>TATTGGAATGATGATCAAGACGCATTTCCATCATAATCAGTTCATCATCGTC |

| | |
|---|---|
| | TCACTCAAACAAGGAATGTTCTCAAATGCAGATGTTCTCTTCCAAACACG TTTTGCCGATGGACATTCTACATGTACACGTCTCAATGGAGGAGATATTGC AGTACTTTGCCAGGATAAAGTTCTACAGGCACAAGCTCTTGAGCTCACAG ATGCTGGAAAAGCAAAGAAGGACGCCGCCGCCAAGAAGGGAGCTCAGA AGAATGACAAAGAA |
| **SMC-4 N-term ATPase Head (codon-optimized for *E. coli*)** | ATGATCCGTAATGTTGAGGTCGATAATTTTAAGTCCTACTTTGGAAAAGCG TCTATCGGCCCATTTCATAAATCCTTTACTTCCATCATTGGACCTAACGGTA GTGGCAAATCAAATCTGATTGACAGCTTGTTGTTTGTTTTCGGCTTCCGTG CCAGTAAGATCCGCAGTGCAAAGGTTAGCAATTTGATTCACAAATCAGCA GGACGTAACCCAGATAAGTGTACAGTGACGATCCACTTCCAACGTATCGTC GATATCCCCGGACATTATGAGGTCGTTAAGGACAGCGAATTTACCATCTCT CGCACTGCTTTTCAGAATAATTCTTCCTCGTATGCTATCGATGGCCGTCCAG CCACGAAGAACGAAGTGGAAGCGCGTCTTCGCCGTGTGGATATTGATATT GAACATAACCGCTTCCTTATTCTTCAGGGCGAGGTCGAGCAAATTGCCATG ATGAAGCCGGTCAAAACGACTAAATCAGAAACGGGGATGGTGGAGTACTT GGAGGACATCATTGGTACTAATCGCTTGGAGCCTTTTGTGAAACTTTTCCA ACGCCGC |
| **SMC-4 C-term ATPase Head (codon-optimized for *E. coli*)** | ATGCGCCTTGAAGAATTCCACAGTGCATTTGAGTTTATCGGCAAGCATTTG GTGGCAGTATTTAAAATGTTGACTGATGGTGGGGACGCCAAGCTGGAGTA CATCGATAAAGACGATCCGTTTCGCCAAGGTATTAGTTTTATGGTTCGTCCA GCGAAGAAAGCTTGGAAGCAGATCCAGTTTCTGAGCGGCGGTGAGAAGA CTTTAAGTAGCTTGGCTTTAATTTTCGCGCTGCATATGTTTCGTCCAACTCC ATTCTACGTGATGGATGAAATCGATGCGGCTTTAGATTACCGTAATGTATCA ATTATCGCTCAATATGTGCGCCAGAAAACTGAGAATGCTCAATTCATTATTA TCTCGTTACGTAACAACATGTTCGAGTTGGCTAATCGCCTGGTTGGCATCT ATAAGGTAGACGGTTGCACCCGTAATGTAGCGATTGATCCTCTTCGTGTTT GCGAGATGGCAAAACAAATCACGGACAGTCTGGGTCAGGCAACGTGTAC GTTACCCGATGAGGTTACACAGCGTTTTAACGAGACTATGTCGCGCCAAA ATAAGGAGATGATCGCTCAAGAA |
| **DPY-27 N-term Forward (BKC.5)** | ATCGGATCCATGATAATTCTGAATATCTA |
| **DPY-27 N-term Reverse (BKC.6)** | ATCGTCGACTTAATGCATTAGTTTGGAAATT |
| **DPY-27 C-term Forward (BKC.7)*** | ATCCATATGGCGAGATTCAACGAGTT |
| **DPY-27 C-term Reverse (BKC.8)*** | ATCCTCGAGTTACTTCTCGCGCCGTTTGAT |
| **MIX-1 N-term Forward (BKC.12)** | ATCCATATGCACATTAAATCCATCCA |
| **MIX-1 N-term Reverse (BKC.14)** | ATCCTCGAGTTACTTCTGATCATACATCTT |
| **MIX-1 C-term Forward (BKC.10)** | ATCGGATCCAAAGTGGACGAGTTGATT |
| **MIX-1 C-term Reverse (BKC.11)** | ATCGTCGACTTATTCTTTGTCATTCTTCTGA |
| **SMC-4 N-term Forward (BKC.25)** | ATCGGATCCATGATCCGTAATGTTGAGGT |
| **SMC-4 N-term Reverse (BKC.26)** | ATCGTCGACTTAGCGGCGTTGGAAAAGTTTCA |
| **SMC-4 C-term Forward (BKC.23)**** | ATCGATATCATGCGCCTTGAAGAATTCCA |
| **SMC-4 C-term Reverse (BKC.24)**** | ATCCTCGAGTTATTCTTGAGCGATCATCTCCT |
| ***dpy-27 (cld21)* repair template (DSgc_006)** | TGCAATGCACCACTACCGTCCAACACCCCTCTACGTGATGGATCAAATCGA TGCAGCACTGGACTTGAACAATGTCAGCCTGATTGCAAACTATATCAAG |

| | |
|---|---|
| ***dpy-27 (cld21)* crRNA (DSgc_g003)** | AGAGCUGAAAGAUCGUUUCA |
| ***dpy-27 E1275Q* screening Forward (BKC.3)** | AGTTCAGTGTACGTCC |
| ***dpy-27 E1275Q* screening Forward (BKC.4)** | ATCGGATCCACCATAATGT |
| ***dpy-27 Indel* Screening Forward (BKC.1)** | TATCACCAATGGCGGAGAT |
| ***dpy-27 Indel* Screening Reverse (BKC.2)** | ATTGCTGAGCCCATTACT |
| **FLAG::*dpy-21* repair template (DSgc_001)** | GTCTGAAAGTGATATAAAATATGAGAAGCACCACTGATTATAAAGACCACGATGGAGACTATAAAGATCATGACATTGACTACAAGGATGACGACGACAAGCTTGCCGCGAATTCGGAGTTTGATAAGGTTTGTTTCGAAGGAAAAAAAATCCC |
| **FLAG::*dpy-21* crRNA (dpy-21)** | GAGAAGCACCACUUUUGAUA |
| **FLAG::*dpy-21* screening Forward (JJ198)** | TTCCGCATTTCTTCTCCGAT |
| **FLAG::*dpy-21* screening Reverse (JJ199)** | TTGGGTGTGCAAGTTGCTCA |

* Both wildtype and EQ mutation gBlocks of DPY-27 C-term were amplified with these primers.

** Primers were used to generate dsDNA for in vitro ATPase reaction (Figure 1 and Supplemental Figure 2)

### Supplemental Table 3 Worm Strains used in this study

| Worm Strain | Source |
|---|---|
| **N2** | **CGC** |
| **EKM 235 *dpy-27 (cld21)* III/qC1 [*dpy-19(e1259) glp-1(q339) qIs26*] III** | **This study** |
| **EKM 190 dpy-27 (cld22) III/qC1 [dpy-19(e1259) glp-1(q339) qIs26] III** | **This study** |
| **EKM40 dpy-28 (tm3535) III/hT2 [bli-4(e937) let-?(q782) qIs48]** | **[90]** |
| **EKM 157 dpy-21(cld12[FLAG::DPY-21]) V** | **This study** |
| **EKM 204 dpy-27 (cld21) III/qC1 [dpy-19(e1259) glp-1(q339) qIs26] III; dpy-21(cld12[FLAG::DPY-21]) V,** | **This study** |
| **EKM 211 dpy-27 (cld22) III/qC1 [dpy-19(e1259) glp-1(q339) qIs26] III; dpy-21(cld12[FLAG::DPY-21]) V** | **This study** |
| **EKM84 unc-119(ed3) III; cldIs12[sdc2::TY1::EGFP::FLAG + unc-119(+))** | **[132]** |
| **EKM272 dpy-27 (cld21) III/qC1[dpy-19(e1259) glp-1(q339) qIs26] III; cldIs12 [sdc2::TY1::EGFP:FLAG + unc-119(+))** | **This study** |
| **EKM275 dpy-27 (cld21) III /qC1 [dpy-19(e1259) glp-1(q339) qIs26] III; cldIs12 [sdc2::TY1::EGFP:3XFLAG + unc-119(+)]; sdc-2(y74) X** | **This study** |
| **EKM268 cldIs12[sdc2::TY1::EGFP::3XFLAG + unc-119(+)]; sdc-2(y74) X** | **This study** |
| **KW 1432 him-8 (e1489) mIs10** | **CGC** |

**Supplemental Table 4 ATP ATPase Assay p-values.** The line indicates the end of comparisons for Figure 1, and that remaining comparisons are for Supplemental Figure 1.

| Šídák's multiple comparisons test | Significance | Adjusted P Value |
|---|---|---|
| **no protein vs. MIX-1 + DPY-27 WT** | **** | <0.0001 |
| **no protein vs. MIX-1 + EQ DPY-27** | ** | 0.0017 |
| **no protein vs. MIX-1 + SMC-4** | **** | <0.0001 |
| **MIX-1 + DPY-27 WT vs. MIX-1 + SMC-4** | *ns* | *>0.9999* |
| **MIX-1 + EQ DPY-27 vs. MIX-1 + SMC-4** | **** | <0.0001 |
| **MIX-1 + DPY-27 WT vs. MIX-1 + EQ DPY-27** | **** | <0.0001 |
| **no protein vs. DPY-27 WT** | **** | <0.0001 |
| **no protein vs. MIX-1** | ** | 0.007 |
| **DPY-27 WT vs. MIX-1** | **** | <0.0001 |
| **MIX-1 vs. MIX-1 + DPY-27 WT** | **** | <0.0001 |
| **DPY-27 WT vs. MIX-1 + DPY-27 WT** | ** | 0.0012 |
| **DPY-27 WT vs. EQ DPY-27** | **** | <0.0001 |
| **DPY-27 WT vs. SMC-4** | ** | 0.0052 |
| **DPY-27 WT vs. MIX-1 + SMC-4** | ** | 0.0011 |
| **MIX-1 vs. EQ DPY-27** | *ns* | *0.5019* |
| **EQ DPY-27 vs. MIX-1 + EQ DPY-27** | ** | 0.0012 |
| **SMC-4 vs. MIX-1 + SMC-4** | **** | <0.0001 |
| **MIX-1 vs. SMC-4** | **** | <0.0001 |
| **EQ DPY-27 vs. SMC-4** | **** | <0.0001 |

**Supplemental Table 5 P-values for Figure 3B.**

| Comparisons | Brood Size | | Embryonic Viability | | Progeny Viability | |
|---|---|---|---|---|---|---|
| **Wildtype vs. *dpy-27 EQ*** | 0.0195 | * | 0.0196 | * | <0.0001 | **** |
| **Wildtype vs. *dpy-27 Indel*** | 0.0168 | * | *0.0825* | *ns* | <0.0001 | **** |
| **Wildtype vs. *dpy-27 EQ/+*** | *>0.9999* | *ns* | *0.3312* | *ns* | *>0.9999* | *ns* |
| **Wildtype vs. *dpy-27 Indel/+*** | *0.9947* | *ns* | *0.136* | *ns* | 0.017 | * |
| ***dpy-27 EQ* vs. *dpy-27 Indel*** | *>0.9999* | *ns* | *0.3456* | *ns* | 0.028 | * |
| ***dpy-27 EQ* vs. *dpy-27 EQ/+*** | 0.0346 | * | *0.2226* | *ns* | <0.0001 | **** |
| ***dpy-27 EQ* vs. *dpy-27 Indel/+*** | 0.0221 | * | 0.04 | * | <0.0001 | **** |
| ***dpy-27 Indel* vs. *dpy-27 EQ/+*** | 0.0013 | ** | *0.3293* | *ns* | <0.0001 | **** |
| ***dpy-27 Indel* vs. *dpy-27 Indel/+*** | 0.0032 | ** | *0.1301* | *ns* | <0.0001 | **** |
| ***dpy-27 EQ/+* vs. *dpy-27 Indel/+*** | *0.9984* | *ns* | *0.7576* | *ns* | *0.4482* | *ns* |

**Supplemental Table 6 P-values for Supplementary Figure 4C.**

| Dunnett's T3 multiple comparisons test | Significance | Adjusted P values |
|---|---|---|
| **Wildtype vs *"WT" dpy-27 EQ/+*** | *ns* | *0.6387* |
| **Wildtype vs *"Mutant" dpy-27 EQ/+*** | **** | <0.0001 |
| **Wildtype vs *dpy-27 Indel/+*** | *ns* | *>0.9999* |
| ***"WT" dpy-27 EQ/+* vs *"Mutant" dpy-27 EQ/+*** | **** | <0.0001 |
| ***"WT" dpy-27 EQ/+* vs *dpy-27 Indel/+*** | *ns* | *0.5703* |
| ***"Mutant" dpy-27 EQ/+* vs *dpy-27 Indel/+*** | **** | <0.0001 |
